# graded-CRISPRi, a novel tool for tuning the strengths of CRISPRi-mediated knockdowns in *Vibrio natriegens* using gRNA libraries

**DOI:** 10.1101/2024.01.29.577714

**Authors:** Daniel Stukenberg, Anna Faber, Anke Becker

**Affiliations:** Center for Synthetic Microbiology, Philipps-Universität Marburg, Marburg, Germany; Department of Biology, Philipps-Universität Marburg, Marburg, Germany

**Keywords:** CRISPR interference, gene repression, gradual downregulation of gene expression, gRNA variants, *Vibrio natriegens*

## Abstract

In recent years, the fast-growing bacterium *Vibrio natriegens* has gained increasing attention as it has the potential to become a next-generation chassis for Synthetic Biology. A wide range of genetic parts and genome engineering methods have already been developed. However, there is still a need for a well-characterized tool to effectively and gradually reduce the expression level of native genes. To bridge this gap, we created graded-CRISPRi, a system utilizing gRNA variants that lead to varying levels of repression strength. By incorporating multiple gRNA sequences into our design, we successfully extended this concept to simultaneously repress four distinct reporter genes. Furthermore, we demonstrated the capability of using graded-CRISPRi to target native genes, thereby examining the effect of various knockdown levels on growth.

## Introduction

*Vibrio natriegens* is the fastest growing organism known to date with a reported doubling time of less than 10 minutes under optimal conditions (Eagon, 1961; Hoffart *et al*., 2017). In addition to its rapid growth, *V. natriegens’* broad substrate spectrum (Hoffart *et al*., 2017) and it’s active metabolism (Long *et al*., 2017; Coppens *et al*., 2023) are considered beneficial properties for biotechnological applications (Hoff *et al*., 2020; Thoma and Blombach, 2021). Furthermore, several pioneering studies have demonstrated the applicability of employing *V. natriegens* for the production of chemical compounds (Dalia *et al*., 2017; Wang *et al*., 2020; Thoma *et al*., 2022). To fully establish *V. natriegens* as a novel, fast-growing model organism, a set of reliable and efficient genetic tools is indispensable. Over the last years, we and others have made significant progress in developing *V. natriegens* into a next-generation SynBio chassis by characterizing standardized genetic parts (Tschirhart *et al*., 2019; Stukenberg *et al*., 2021; Tietze *et al*., 2022) and developing genome engineering tools (Weinstock *et al*., 2016; Dalia *et al*., 2017; Stukenberg *et al*., 2022). These tools allow the construction of heterologous pathways or genetic circuits, as well as the introduction of permanent genetic modification into the genome of *V. natriegens*.

However, a set of tools for the precise regulation of native gene expression in *V. natriegens* is not yet available. In particular, a system enabling graded repression of gene expression, as opposed to a binary on/off switch, would support or even enable a broad spectrum of basic and applied scientific research in this chassis. Among various possibilities, such an approach could lead to interesting insights into the metabolism of *V. natriegens*. For example, recent studies in *Escherichia coli* showed that the abundance of many proteins can be reduced with little to no effect on growth, even though the respective genes are essential under the specific conditions (O’Brien, Utrilla and Palsson, 2016; Sander *et al*., 2019; Hawkins *et al*., 2020). From an application-oriented perspective, it has been demonstrated that repression of competing pathways, resulting in redirection of metabolic fluxes, can enhance the efficiency of a heterologous pathway. (Kim *et al*., 2017; Tian *et al*., 2019).

To enable similar approaches in *V. natriegens*, we developed a CRISPRi-based tool for the targeted repression of endogenous genes. CRISPRi uses a catalytically inactive Cas9 (dCas9), which is guided to a DNA sequence by a programmable gRNA to block transcription (Qi *et al*., 2013). CRISPRi was used before in *V. natriegens* to perform a genome-wide screen identifying essential genes and those necessary for rapid growth under specific conditions (Lee *et al*., 2019). In the referenced study, the CRISPRi tool lacked tight control, and the authors took advantage of the system’s inherent leaky expression to generate growth-defective phenotypes (Lee et al., 2019). However, the leakiness of this system complicates the study of essential genes as it creates a strong selection pressure for adaptational mutations or inactivation of the CRISPRi system itself. In our study, we therefore envisioned a CRISPRi-based tool that can generate libraries of strains with different knockdown strengths. This approach is especially beneficial in scenarios where the level of repression required to achieve a desired phenotype is lacking.

Here, we report the development and characterization of graded-CRISPRi as a novel tool for *V. natriegens*. By using two different inducible promoters, we achieved tight regulation of the CRISPRi system that did not show a measurable effect in the absence of inducers. Next, we achieved graded repression strengths by varying the binding strength of the dCas9-gRNA complex to the DNA target sequence. This was achieved by using gRNA libraries with spacers of different lengths and with various mismatches to the target sequence. We further demonstrated the simultaneous targeting of multiple reporter genes with several gRNAs, leading to a set of strains with different expression levels of each reporter gene. Lastly, we used graded-CRISPRi for the down-regulation of metabolic genes in *V. natriegens* and investigated the phenotype resulting from different knockdown strengths.

## Results and discussion

### Establishing a tightly controlled CRISPRi tool for *V. natriegens*

As a first step in the development of graded-CRISPRi, we focused on the tight regulation of CRISPRi in *V. natriegens*. It was shown previously that the activity of CRISPRi-based tools is dependent on sufficient expression of both *dcas9* and *gRNA* (Fontana *et al*., 2018). Thus, we concluded that employing two orthogonal inducible promoters to control the expression of both genes should achieve the most stringent control of the system. This is because simultaneous leaky expression of *dcas9* and *gRNA* in the same cell would be necessary to produce a functional dCas9-gRNA complex. A further rationale for utilizing two separate inducible promoters for *dcas9* and *gRNA* expression was the ability to independently regulate the expression strength of both CRISPRi components. This strategy was vital to mitigate toxicity and reduce metabolic burden. For the two inducible promoters, we chose modified versions of the P_tet_ and P_3b5b_ inducible promoters with their corresponding inducers anhydroteracycline (ATc) and dihydroxybenzoic acid (DHBA) (Meyer *et al*., 2019). To benchmark the new inducible CRISPRi system in *V. natriegens*, we targeted a chromosomally integrated and strongly constitutively expressed *mScarlet-I* reporter gene (Fig. 1a).

**Figure 1:**
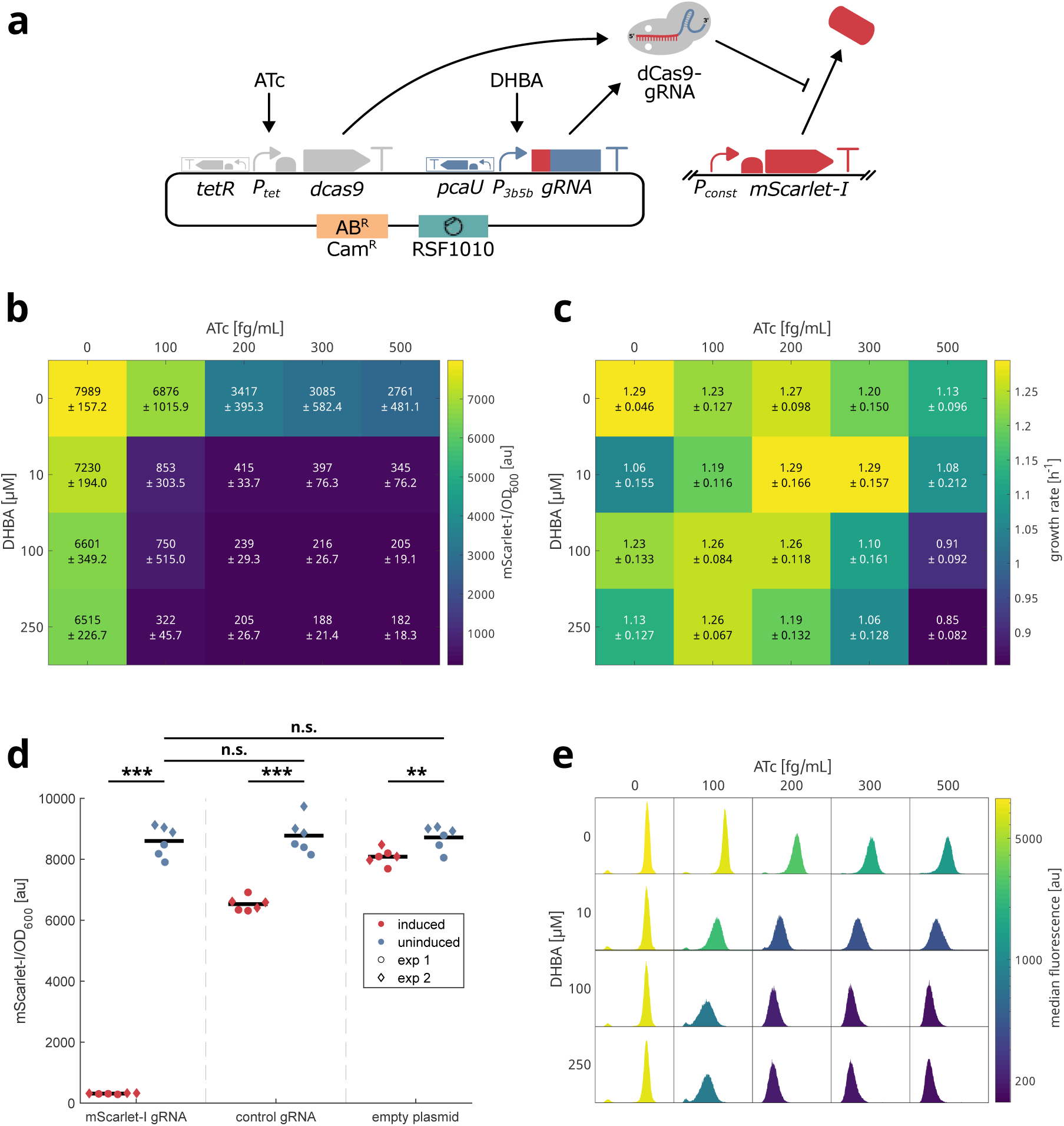
Initial characterization of CRISPRi in *V. natriegens*. **(a)** System for the characterization of the CRISPRi tool in *V. natriegens*. All components of the CRISPRi system were located on a single plasmid (pST_300). Expression of *dcas9* is controlled by a P_tet_ promoter, which can be activated through addition of ATc. Expression of the gRNA is controlled through a P_3b5b_ promoter and is activated by DHBA. The gRNA is composed of a target specific spacer sequence (red) and a gRNA scaffold sequence (blue). This system is tested in a strain carrying a chromosomally integrated mScarlet-I expression cassette (DST018). Expression of mScarlet-I is controlled through a strong constitutive promoter (P_J23111_). **(b, c)** Testing combinations of DHBA and ATc for their impact on knockdown strengths **(b)** and growth rate **(c)**. Colors in the heatmap indicates growth rate of the cultures. Embedded text reports the mean growth rate, as well as the standard deviation from the mean. This data is based on two independent experiments with three biological replicates. Experiments were performed in M9G. **(d)** Testing inducibility of the CRISPRi system and comparison with control constructs. Induced samples (200 fg/mL ATc, 100 µM DHBA) and uninduced samples are shown in red and blue, respectively. Data points from the first independent experiment are shown as circles and data from the second experiment are displayed as diamonds. Control gRNA refers to a gRNA targeting *dns*, which is deleted in this strain. The experiment was performed in M9G with CRISPRi plasmid pST_300 in DST018. Significances were calculated with a two-sample t-test. n.s.: p > 0.05, *: p > 0.01, **: p > 0.001, ***: p < 0.001. **(e)** Analyzing mScarlet-I signal resulting from different combinations of DHBA and ATc at the single cell level. Fluorescence was measured with a flow cytometer. Displayed is a representative histogram from six measurements (three biological replicates, and two independent experiments). The color of the histograms represents the median fluorescence. Samples were drawn from cultures grown in M9G with CRISPRi plasmid pST_300 in strain DST018.

Firstly, we measured the mScarlet-I signal from *V. natriegens* cultures in minimal medium (M9 + Glucose = M9G) and rich medium (LBv2) with different inducer concentrations. In M9G, we achieved an approximately 40-fold reduction of the mScarlet-I signal at the highest tested concentrations of ATc and DHBA. As expected, this strong effect was only visible when both inducers were added (Fig. 1b). However, induction of *dcas9* with ATc alone also resulted in a three-fold repression, while induction of *gRNA* with DHBA alone led to a smaller but still noticeable decrease in reporter signal (1.25-fold) (Fig. 1b). When comparing growth rates resulting from cells treated with different inducer concentrations, it became apparent that especially the addition of ATc resulted in a noticeable decrease of up to 1.2-fold, at the highest tested concentration (Fig. 1c). This is most likely the consequence of a metabolic burden, which we described before for high expression of reporter genes (Stukenberg *et al*., 2021). Alternatively, unspecific toxicity of dCas9 (Cho *et al*., 2018) could be an explanation for the decrease in growth rate. We finally decided to use 200 fg/mL for ATc and 100 µM DHBA for further experiments in M9G, as we considered these concentrations as a good middle ground between strong gene repression and an acceptable effect on growth.

We also determined optimal inducer concentrations for application of this CRISPRi system in LBv2. Similar to the results from experiments in M9G, only addition of both inducers led to the strongest mScarlet-I repression. Interestingly, we only achieved an approximately 8-fold reduction of the mScarlet-I signal in LBv2, compared to a 40-fold reduction in M9G. Additionally, a higher concentration of ATc was required to achieve this repression (Fig. S1). Unexpectedly, we observed a slightly increased growth rate at low inducer concentration in LBv2, possibly because the reduced expression of *mScarlet-I* overcompensated the burden arising from expression of *dcas9* (Fig. S2). For further experiments in LBv2, we decided to use 1000 fg/mL for ATc and 100 µM DHBA. Overall, differences in growth rate in LBv2 were smaller than in M9G (Fig. S2, Fig. 1c).

To evaluate the leakiness of the inducible CRISPRi system, we compared the mScarlet-I signals across three strains, each carrying a different plasmid. The first strain was equipped with a CRISPRi system targeting mScarlet-I, the second with a CRISPRi system featuring a non-binding gRNA, and the third carried an empty plasmid devoid of any CRISPRi system. Neither in M9G (Fig. 1d) nor in LBv2 (Fig. S3), a significant difference was detected between the strains confirming a tight regulation of the graded-CRISPRi system. However, we observed that induction of the CRISPRi system with the non-binding control gRNA in M9G resulted in an unexpected drop of reporter signal (1.3-fold) (Fig. 1d). To investigate whether this was caused by expression of *dcas9*, *gRNA* or both, we measured mScarlet-I signal at different concentrations of both inducers in cells carrying a CRISPRi system encoding the non-binding control gRNA construct. We found that the combination of both inducers led to the strongest reduction of mScarlet-I signal (Fig. S4). Although determining the exact mechanism behind this effect is beyond the scope of this work, we assume that a regulatory response of the cells reduces transcription from the constitutive promoter of *mScarlet-I*. It was shown before in *E. coli* that production of heterologous proteins in *E. coli* can trigger the σ^32^-mediated heat shock response, which reduces expression from σ^70^-dependent constitutive promoters through sigma factor competition (Ceroni *et al*., 2018). Sequestration of RNA polymerases for the transcription of *dcas9* and *gRNA* is likely to also result in a reduction of free RNA polymerases.

One desired feature of the graded-CRISPRi tool is the generation of libraries of variants leading to different knockdown strengths. We saw that different inducer concentrations lead to different reporter signals and therefore to different knockdown strengths. However, previous studies have shown that controlling CRISPRi repression strength through limiting either dCas9 or gRNA availability results in high population heterogeneity (Vigouroux *et al*., 2018). To determine if this is also the case with our system in *V. natriegens*, we measured the mScarlet-I signal at the single cell level in a flow cytometer. At all concentrations, we observe a unimodal distribution (Fig. 1e). However, consistent with literature on *E. coli* (Vigouroux *et al*., 2018), the histogram at moderate inducer concentrations is lower but wider compared to higher concentrations, indicating a high population heterogeneity both in M9G (Fig. 1e) and LBv2 (Fig. S5). Consequently, we did not consider adjusting CRISPRi strength through titration of dCas9 and gRNA expression to be a suitable strategy to obtain graded knockdown levels in *V. natriegens*.

### Achieving a graded-CRISPRi repression through gRNA libraries

As a promising alternative to tuning inducer concentrations, we considered using gRNAs with mismatches to the target sequence to adjust CRISPRi knockdown efficiency, similar to previous work (Hawkins *et al*., 2020). As a proof of concept, we exemplarily tested the targeting of mScarlet-I using a gRNA sequence with two defined mismatches at position five and ten from the PAM proximal nucleotide (C5T, A10G). As a result, we obtained an approximately seven- and two-fold reduction of the mScarlet-I signal in M9G and LBv2, respectively (Fig. 2a, Fig. S6). Hence, we found a weaker repression of the mScarlet-I signal in both minimal and rich media in comparison to a CRISPRi system with a perfectly matching gRNA.

**Figure 2:**
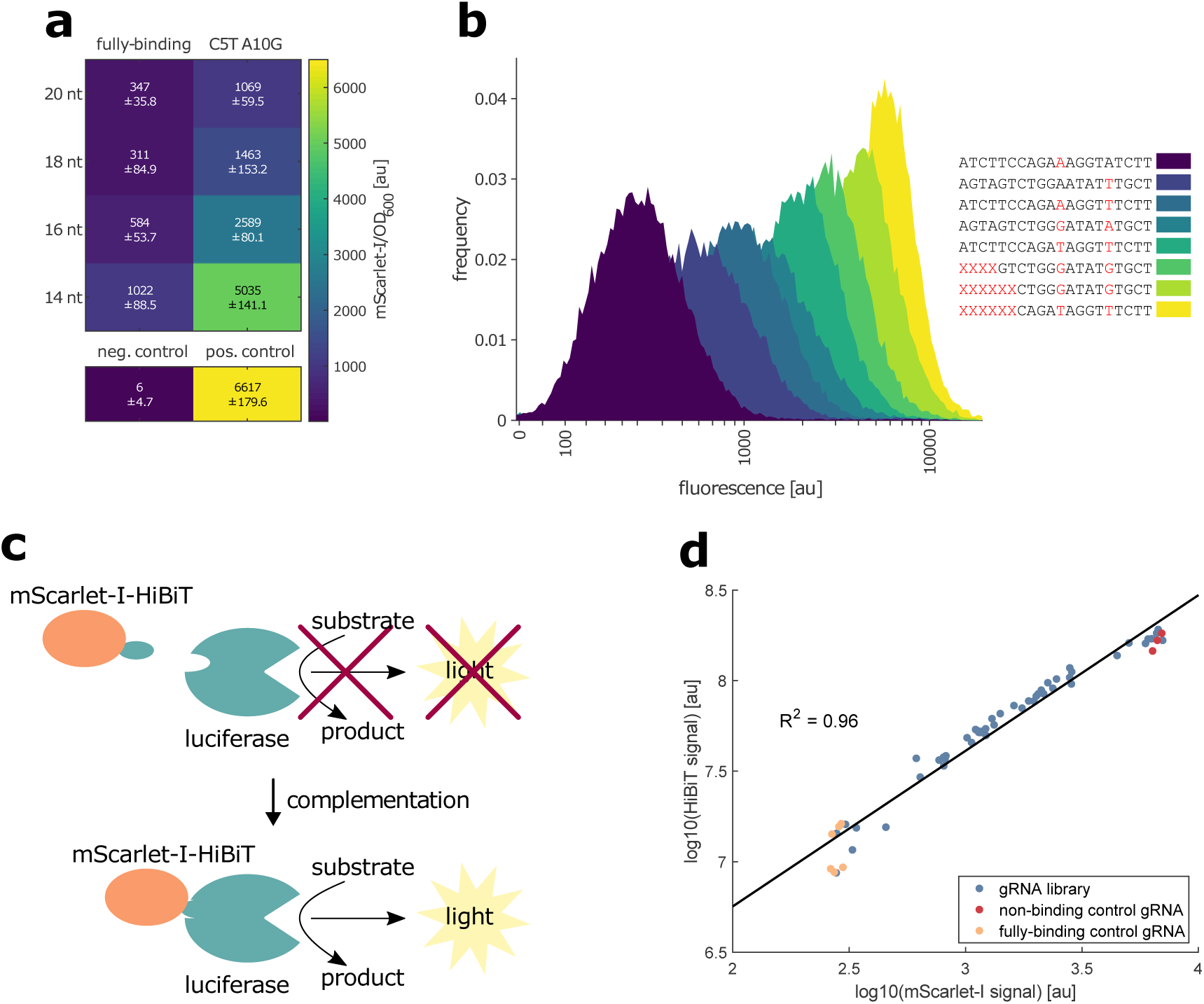
Testing mismatched and truncated gRNAs for CRISPRi in *V. natriegens*. **(a)** Analyzing repression strengths of gRNAs with mismatches and different spacer lengths. C5T and A10G indicate mismatches to the target sequence and are counted form the PAM proximal nucleotide of the spacer sequence. neg. control: strain without integrated mScarlet-I cassette (DST016). pos. control: strain integrated mScarlet-I cassette (DST018) with a CRISPRi plasmid with a non-binding control gRNA. Colors in the heatmap indicate the mScarlet-I fluorescence signal. Embedded text represents the mean mScarlet-I signal, as well as the standard deviation from the mean. This data is based on two independent experiments with three biological replicates. Experiments were performed in M9G with the CRISPRi plasmid pST_300. **(b)** Analyzing fluorescence from gRNA variants by flow cytometry. Histograms from selected variants are displayed and their gRNA spacer sequence is provided in the legend. Coloring does not reflect fluorescence and only serves to distinguish samples. Red letters in the legend indicate mismatches in the gRNA variants. Red “X” letters indicate truncations. **(c)** Scheme of the HiBiT tag system. The HiBiT tag is attached to the C-terminal end of a protein of interest (here mScarlet-I). This HiBiT tag complements an otherwise unfunctional luciferase core enzyme, leading to the generation of light. **(d)** Correlation between mScarlet-I and HiBiT signal. The experiment is based on a gRNA library targeting mScarlet-I (see Fig. 2b) in a strain with a chromosomally integrated mScarlet-I-HiBiT expression cassette (DST018). Each data point represents a single measurement from a variant of the gRNA library (blue), the non-binding control gRNA (red) or the fully-binding gRNA (yellow). R^2^ is derived from a linear regression of the log10 fluorescence and luminescence values. Experiments performed in M9G with induction of the CRISPRi system (200 fg/mL ATC, 100 µM DHBA).

One desired capability of the graded-CRISPRi tool is to generate mild knockdowns to reveal interesting phenotypes at a reduced abundance of essential proteins. We assume that even the reduced repression strength, which we obtained with the exemplarily tested C5T, A10G mismatches, would still be too strong and prevent growth when targeting essential genes. To generate gRNA variants with even weaker DNA binding strengths, we tested truncated gRNA sequences in combination with the above-described mismatches. Truncated gRNAs have been used before to reduce off-target effects of CRISPR-Cas9 systems for genome editing. The reduction in off-target effects results from weakening the DNA binding strength of the Cas9-gRNA complex through gRNA truncation (Fu *et al*., 2014). When using the *mScarlet-I* targeting gRNAs with the spacer sequence truncated to 14 nucleotides featuring C5T and A10G mismatches, the knockdown effect was reduced to just 1.3- and 1.1-fold in M9G and LBv2, respectively (Fig. 2a, Fig. S6). Intermediate knockdown effects were obtained for gRNA variants with shorter truncation or no mismatches (Fig. 2a). These results encouraged us to design libraries of gRNAs with spacers of different lengths (20, 16, and 14 nucleotides) in combination with random mismatches at position five and ten from the PAM-proximal nucleotide. Additionally, we included a second distinct gRNA sequence which targeted a different section of the *mScarlet-I* sequence to expand the range of possible knockdown effects even further.

In theory, 96 distinct gRNA (2 x 2^4^ x 3 = 96) can be obtained by combining two different gRNA sequences with two randomized nucleotides and three different lengths. We designed and constructed such a library targeting *mScarlet-I*. We then introduced this plasmid library into *V. natriegens* and identified the gRNA spacer sequence of 48 randomly picked colonies by Sanger sequencing, eventually obtaining 33 unique gRNA variants (Table S1). To check if this is within the expected range, we calculated a probability distribution for the number of unique variants when 48 variants are randomly selected from a library of 96 possible and equally probable variants. In this distribution, finding 38 unique variants is the most probable outcome, which is slightly higher than the 33 unique variants that we found here (Fig. S7). Nevertheless, we concluded that we generated a sufficiently large sequence diversity to proceed with an experimental characterization of these gRNA variants. Consequently, we next assessed the repression strength of these gRNA variants by analyzing their targeting of *mScarlet-I* at the single-cell level. For this, we employed flow cytometry to resolve differences between individual cells within the population. As expected, we observed very high to very low mScarlet-I signals, indicating weak and strong repression, respectively (Fig. 2b).

Next, we examined whether modifying repression strength through limiting inducer concentrations or by using a CRISPRi system with one of the gRNA library variants leads to a more homogenous population. Therefore, we compared samples yielding a similar median fluorescence signal originating from either strategy. In accordance with literature (Vigouroux *et al*., 2018), our flow cytometry data revealed that strains with a gRNA library variant exhibited a slightly narrower and therefore more uniform fluorescence distribution compared to those with a fully binding gRNA but with reduced inducer concentrations. This finding confirms that the goal of attenuated repression of gene expression can be achieved better with lower cell-to-cell variability by using mismatch gRNA libraries, as opposed to attenuation by lower *gRNA* or *dcas9* induction (Fig. S8).

### Tracking the reduced protein abundance using the HiBiT tag system

So far, the characterization of the graded-CRISPRi tool relied on the simple quantification of mScarlet-I as a fluorescent protein. To apply graded-CRISPRi for the repression of other genes, a convenient strategy is desired to quantify the resulting change in protein abundance. While adding a fluorescent reporter as a tag to the target protein seems to be the most obvious approach, it comes with several drawbacks. Firstly, the fused fluorescent protein would be expressed in a 1:1 stoichiometry with the tagged protein. For this strategy, the inevitable translation of an additional ∼ 250 amino acids for each fluorescent tag could result in a substantial metabolic burden which might significantly impact the strain’s phenotype. These concerns would particularly apply when examining highly abundant proteins and therefore reduce the applicability of this strategy. Secondly, attaching fluorescent tags to proteins for tracking expression levels carries the risk of adversely affecting the functionality of the tagged proteins. Lastly, the limited sensitivity of fluorescent proteins might set limits to the detection of protein abundance, especially for less abundant protein targets.

To mitigate these limitations, we tested the HiBiT peptide tag system, which was developed and commercialized by Promega (Schwinn *et al*., 2018). The HiBiT tag is an eleven amino acids long peptide, which complements a luciferase enzyme to generate bioluminescence (Schwinn *et al*., 2018) (Fig. 2c). The emitted signal is supposed to follow a linear correlation with the abundance of the HiBiT peptide and therefore with the tagged protein of interest. Due to the ∼20-fold shorter size of the HiBiT tag compared to a fluorescent protein, we expect a neglectable metabolic burden caused by the additional synthesis of the tag. Furthermore, we anticipate a reduced risk of impairing the target protein’s functionality due to the smaller size of the HiBiT tag compared to that of a fluorescent protein. These benefits motivated us to use this reporter system for protein quantification in our study. However, it is important to acknowledge that a limitation of the HiBiT tag system remains in its reliance on an endpoint enzyme reaction for signal measurement. Therefore, this approach does not allow for tracking of the change in protein abundance over time, unless samples are taken at several timepoints during the experiment.

To assess the HiBiT system’s effectiveness in quantifying protein abundance, we engineered a C-terminal fusion of mScarlet-I with the HiBiT tag. Consequently, we anticipated a linear correlation between fluorescence and luminescence signals, derived from the 1:1 stoichiometry between the fluorescent reporter mScarlet-I and the attached HiBiT tag. Subsequently, we employed the strain library with gRNA variants targeting mScarlet-I to create a spectrum of protein abundance levels in the different *V. natriegens* strains. As a result, we obtained a strong linear correlation between both reporter signals with R^2^ = 0.96 (Fig. 2d). In a comparable experiment using the firefly luciferase (Fluc) in place of mScarlet-I with the HiBiT tag, we achieved an even stronger correlation, with an R² value of 0.99 (Fig. S9).

### Targeting multiple reporters simultaneously

Furthermore, we wondered if we could expand graded-CRISPRi to target multiple genes simultaneously. Since CRISPR-Cas9-based systems are inherently modular, repression of multiple targets can be accomplished by integrating several *gRNA* expression cassettes into the plasmid that carries the *dcas9* sequence. Our previous experiments have shown that a strong repression of the target gene through the CRISPRi system depends on the sufficient expression of both *gRNA* and *dcas9* (Fig. 1b, Fig. S1). Therefore, we reasoned that we would require stronger *dcas9* induction, as introducing additional gRNA cassettes leads to a higher total gRNA abundance. To test this, we created a set of plasmids encoding one *mScarlet-I* targeting gRNA and up to three additional non-binding decoy gRNAs. Subsequently, we evaluated the repression strength of the mScarlet-I reporter across strains carrying plasmids encoding zero to three decoy gRNAs. As expected, addition of decoy gRNAs reduced the knockdown strength at our usual ATc inducer concentration of 200 fg/mL in M9G. This indicates that dCas9 is not sufficiently abundant to efficiently repress the expression of the target when more than one *gRNA* cassette is present. This effect was not observed at higher ATc concentrations when more dCas9 is produced.

Specifically, a concentration of 600 fg/mL of ATc is necessary with three decoy *gRNA* cassettes to achieve the same level of *mScarlet-I* repression as with constructs without additional gRNA cassettes. (Fig. 3a). Unfortunately, this increased *dcas9* expression also leads to a noticeable reduction in growth rate (Fig. S10). Therefore, it’s important to always consider the trade-off between strong repression and an undesired impact on growth, carefully balancing the implications of either a weaker repression or a more pronounced growth defect.

**Figure 3:**
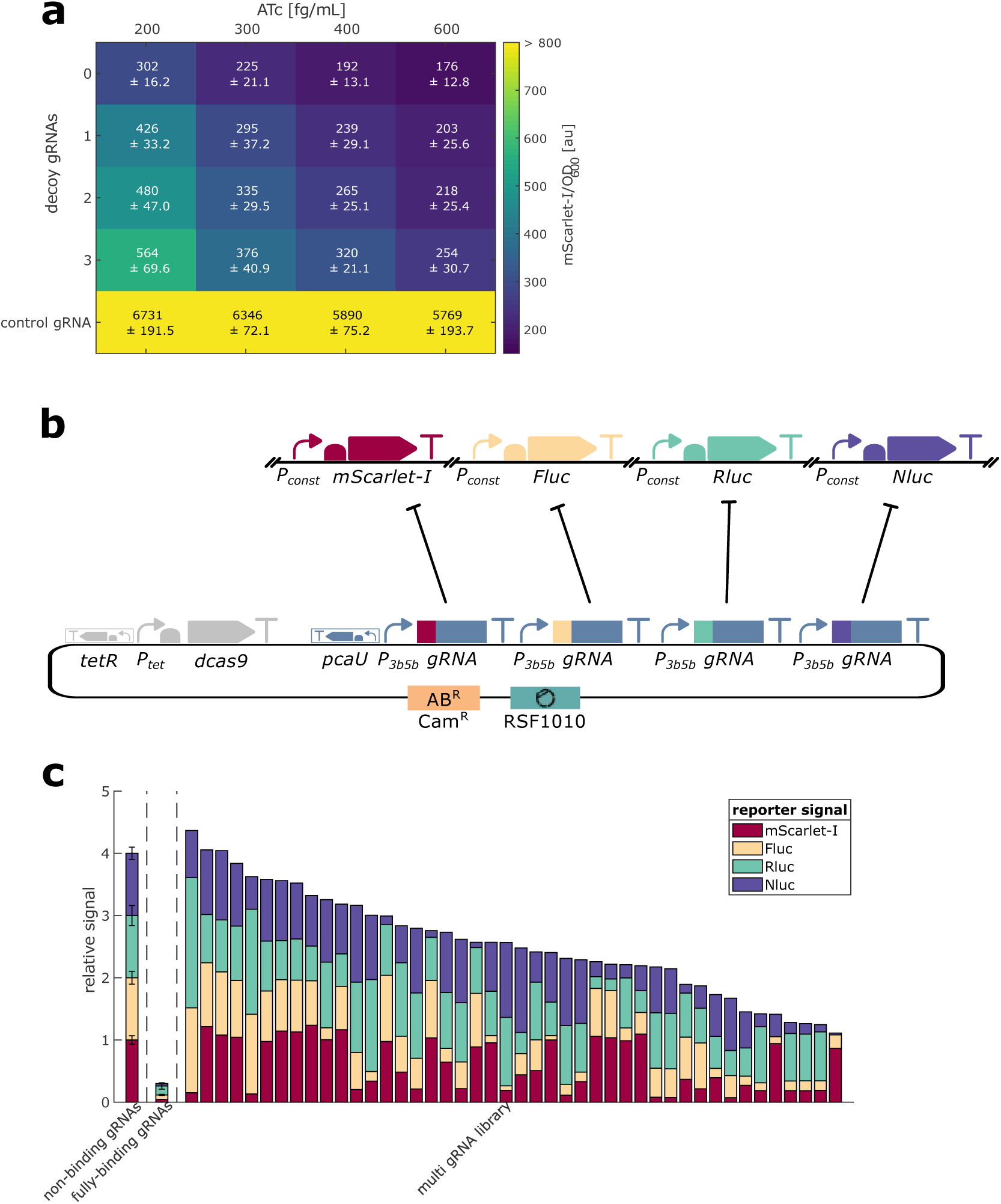
Expanding graded-CRISPRi to multiple targets. **(a)** Testing the effect of additional gRNAs on repression strength at different ATc concentrations. CRISPRi plasmids encoding a gRNA targeting mScarlet-I and up to three decoy gRNAs (targeting *dns* = non-binding gRNA) were tested at different ATc concentrations and the same DHBA concentration (100 µM). Colors in the heatmap indicate mScarlet-I fluorescence signal. Embedded text represents the mean mScarlet-I signal, as well as the standard deviation from the mean. This data is based on two independent experiments with three biological replicates. Experiments were performed in M9G in strain DST018. **(b)** Scheme for the simultaneous targeting of four reporter genes. The reporter genes are integrated at distinct sites in chromosome 1 (mScarlet-I, Fluc, Rluc) and chromosome 2 (Nluc) and are expressed from constitutive promoters. The CRISPRi plasmid encodes four gRNA expression cassettes, each targeting one reporter gene. **(c)** Targeting four reporter genes with gRNA libraries. The controls carry either a CRISPRi plasmid encoding four non-binding gRNAs or one fully-binding gRNA for each reporter. The stacked bars show the relative signal compared to the control with four non-binding gRNAs. Error bars displayed for the controls indicate the standard deviation from the mean of three biological replicates. Stacked bars of the library clones were sorted according to the sum of all relative reporter signals. Experiments were performed in M9G with induction of the CRISPRi system (600 fg/mL ATC, 100 µM DHBA).

We could already show that generating libraries of gRNA sequences with mismatches to the target leads to variable repression strengths. This motivated us to test if we can also apply this approach to multiple targets. To distinguish the repression of multiple targets, we created a *V. natriegens* strain with four different reporter cassettes integrated into distinct regions of the genome. We used mScarlet-I, firefly luciferase (Fluc), renilla luciferase (Rluc), and nano luciferase (Nluc). After creating a *V. natriegens* strain with all four reporters, as well as strains with each reporter alone, we performed an initial experiment to check for crosstalk between the reporters. We observed no crosstalk, with the exception of Nluc, which generated a moderate luminescence signal with Coelenterazine, the substrate of Rluc (Fig. S11). For the goal of this experiment, we consider this moderate crosstalk acceptable but note that the measured value for Rluc might be slightly overestimated due to the reaction of Nluc with the Rluc substrate.

To evaluate graded-CRISPRi against multiple targets, we created a plasmid library, with each variant encoding a different combination of different gRNAs targeting the four reporters (Fig. 3b). Upon testing this library in the strain containing four reporters, we observed a wide range of outcomes, which varied depending on the gRNA variants utilized in the CRISPRi system. Certain combinations of gRNA variants led to strong repression of one or two reporters, while other gRNA applications resulted in moderate repression across all four reporter genes (Fig. 3c). There was no correlation between the four reporter signals, indicating that this approach led to independent repression of all four reporter genes (Fig. S12). Therefore, graded-CRISPRi with multiple gRNAs holds potential for future applications in tuning the expression levels of several endogenous target genes, such as in metabolic engineering projects. This could be used to fine-tune through gene expression control levels of enzymes in pathways either feeding into or competing with an introduced heterologous pathway. Then, variants with the desired phenotype could be screened for, followed by investigating which changes have led to this result.

### Targeting native genes

So far, we exclusively targeted chromosomally integrated reporter genes. While this allowed for the characterization of the graded-CRISPRi tool, targeting native genes is clearly the more relevant application for future research. As a proof of concept, we selected four endogenous metabolic genes of *V. natriegens* – *eno*, *metA*, *metE*, and *argA*. According to a CRISPRi screen conducted by Lee et al. (2019), repressing *eno* and *metA* led to reduced fitness in minimal media, which aligns with their roles in glycolysis and amino acid biosynthesis, respectively. In contrast, targeting *metE* and *argA* appeared to have no significant effect (Lee *et al*., 2019). This outcome was unexpected, especially since a similar CRISPRi screen in *E. coli* (Wang *et al*., 2018) highlighted the importance of *metE* and *argA* for growth in minimal medium. Given this context, we chose to target *eno*, *metA*, *metE*, and *argA* in *V. natriegens* using our graded-CRISPRi tool. Our aim was to explore the effects of varying repression strengths on these genes. Hence, we created translational fusions of *eno*, *metA*, *metE*, and *argA* with the HiBiT tag to enable quantification of the knockdown strengths. Firstly, we could show that the integration of the HiBiT tag did not affect growth of the resulting strains (Fig. S13). Secondly, we created gRNA libraries targeting each of these endogenous genes. For *eno* and *metA*, this resulted in a diverse array of growth phenotypes (Fig. 4a, Fig. 4b). In contrast, all samples targeting *metE* and *argA* exhibited growth patterns similar to strains with the non-binding control gRNA. (Fig. 4c, Fig. 4d). These results are in accordance with the data from a previously published CRISPRi screen in *V. natriegens* (Lee *et al*., 2019).

**Figure 4:**
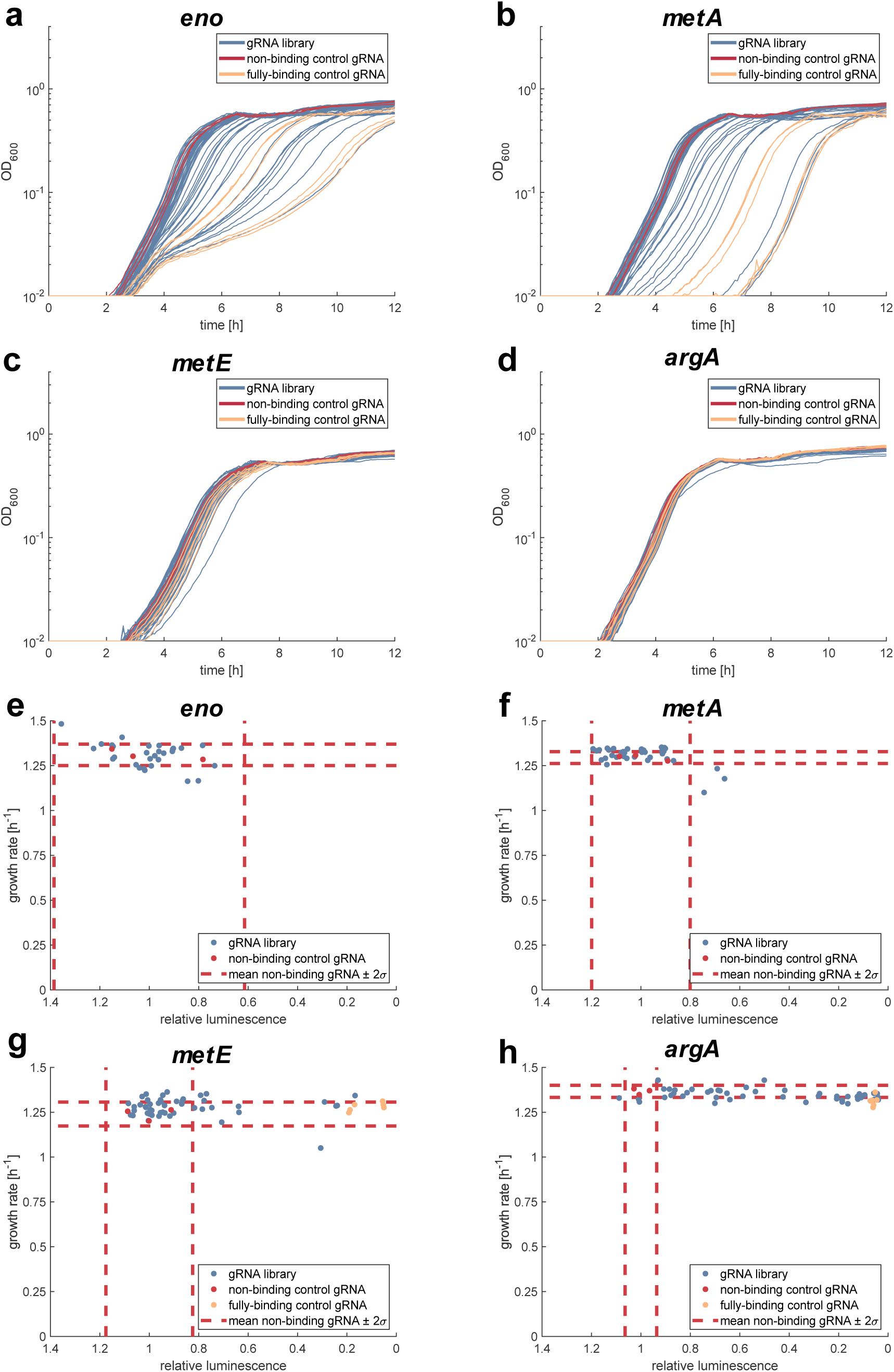
Targeting native genes with graded-CRISPRi. **(a - d)** Growth curves resulting from targeting of native genes by graded-CRISPRi. Each line represents one variant of the gRNA library (blue), the non-binding control gRNA (red) or the fully-binding gRNA (yellow). Strains carry a C-terminal HiBiT tag fusion of the target protein **(e - h)** Scatter plot between growth rate and luminescence from the HiBiT tag. Relative luminescence was calculated relative to the mean of the values obtained from the non-binding control strains. Each data point represents a single measurement from a variant of the gRNA library (blue), the non-binding control gRNA (red) or the fully-binding gRNA (yellow). Dotted red lines indicate the mean values from three replicates of the non-binding control gRNA plus and minus two standard deviations. Only samples with a growth behavior similar to the non-binding control strains were analyzed. Experiments were performed in M9G with induction of the CRISPRi system (200 fg/mL ATC, 100 µM DHBA). The CRISPRi plasmid pST_300 was used as well as the strains DST025 (eno- HiBiT), DST024 (metA-HiBiT), DST031 (metE-HiBiT), DST023 (argA-HiBiT).

Moreover, it was shown before in *E. coli* that some important enzymes are overabundant and that their abundance can be significantly reduced without an effect on growth (Sander *et al*., 2019). We therefore wanted to test if we can find variants in the *eno* and *metA* libraries with reduced abundance of the respective protein but without an effect on growth. For example, if we observe a strain with a two-fold reduced enzyme abundance but with a control-like growth rate, this will tell us that the respective enzyme has a higher abundance than necessary to sustain fast growth.

To quantify the abundance of the chosen enzymes, we repeated the cultivation of the samples without a growth defect and harvested the cultures in the exponential phase. We then proceeded with the quantification of the HiBiT tag to investigate the abundance of the tagged enzymes. Fig. 4e – Fig. 4h illustrate the relationship between growth rate and luminescence signal. For *metA* and *eno*, almost all included samples are within two standard deviations of the values for the non-binding control gRNA (Fig. 4e, Fig. 4f). None of the strains with a control-like growth rate had a significantly reduced luminescence signal. The absence of strains with a control-like growth behavior but with reduced luminescence suggests that even a moderate reduction of the respective enzymes affects growth. The picture is different for *metE* and *argA* (Fig. 4g, Fig. 4h). We found variants with 10 to 20-fold reduced luminescence signals but without an effect on growth rate. This confirms that our CRISPRi tool can lead to a strong repression of native genes. Also, this result is in accordance with the results from the CRISPRi screen (Lee *et al*., 2019), which showed no effect of a CRISPRi mediated knockdown of *metA* and *argA* on growth in minimal medium. However, even with a 10 to 20-fold reduction in abundance, it cannot be excluded that the enzyme level is still sufficient to support growth. We therefore created deletion strains of *argA* and *metE* and tested their ability to grow in M9G. Those strains did not show a major grow defect compared to the parental strain, confirming that these genes are not necessary for growth of *V. natriegens* in M9G (Fig. S14). Literature provides some insights into these observations. Several marine bacteria possess an *argH(A)* gene encoding a bifunctional enzyme capable of catalysing both the initial and final steps of arginine biosynthesis (Xu *et al*., 2000, 2006). This could compensate for the loss of *argA* in our gene deletion experiment and explain consistent growth rates. The larger size of ArgH in *V. natriegens* compared to *E. coli* (624 vs. 456 amino acids) suggests that *V. natriegens*’ ArgH might also function as a bifunctional ArgH(A) enzyme, functionally replacing ArgA. Regarding *metE*, the presence of a second ORF (PN96_19000) with the same annotation might indicate a gene duplicate, which we speculate could compensate for *metE* loss.

## Conclusion

We developed and characterized a tightly controllable CRISPRi system for *V. natriegens*. By using libraries of gRNA sequences leading to a weaker binding of the dCas9-gRNA complex to the target DNA sequence, we obtained graded repression. We could show that graded-CRISPRi can be used to target native genes, which resulted in diverse growth behaviors. In future studies, this approach could be expanded to target other essential or non-essential genes to study the phenotypes resulting from different knockdown strengths. To allow quantification of the resulting protein abundance, we established the HiBiT tag system for use in *V. natriegens*. Beyond being used in combination with graded-CRISPRi, the HiBiT tag system might be an interesting alternative for the quantification of the abundance of single proteins in different strains or different conditions. We demonstrated the applicability of graded-CRISPRi for the simultaneous repression of four reporter genes. This could be used in studies where only a specific combination of expression levels from multiple genes leads to a favorable outcome.

## Methods

### Bacterial strains and culture conditions

All strains created in this work are based on a *V. natriegens* strain ATCC14048 with a deletion of the *dns* gene (Stukenberg *et al*., 2021). This gene codes for a DNA endonuclease, which prevents efficient transformation with plasmid DNA using the heat-shock transformation protocol. *V. natriegens* was routinely grown in LBv2 (Weinstock *et al*., 2016). Whenever selection was required, chloramphenicol was added to the medium with a final concentration of 4 µg/mL or 2 µg/mL for liquid and solid medium, respectively. Glycerol stocks were prepared for long term storage at -80°C by growing cultures for 6 – 8 h at 37 °C and mixing 700 µL of the grown cultures with 700 µL of 50% glycerol. Experiments were performed either in buffered M9 minimal medium (recipe in Tables S2 – S5) with glucose as the sole carbon source (M9G) or in LBv2 (Weinstock *et al*., 2016).

### Preparation of chemically competent *V. natriegens* cells and heat-shock transformation

Preparation of chemically competent cells and heat-shock transformation were performed as described before (Stukenberg *et al*., 2021). A preculture of the respective *V. natriegens* strain was inoculated from a glycerol stock and grown overnight at 30 °C and 200 rpm. On the next day, 125 mL of preheated LBv2 medium (37 °C) were inoculated with the overnight culture to a final OD_600_ of 0.01 in a 1 L baffled shake flask. This culture was grown at 200 rpm (37 °C) until an OD_600_ between 0.5 and 0.7 was reached. The culture was then transferred to pre-chilled 50 mL falcon tubes and incubated on ice for 10 min, followed by centrifugation for 10 min at 3000 × g at 4 °C. The supernatant was discarded, and the pellets were resuspended in 40 mL cold TB buffer per 125 mL bacterial culture (TB buffer: 10 mM Pipes, 15 mM CaCl2, 250 mM KCl, pH adjusted to 6.7 with KOH, then add 55 mM MnCl2, filter sterilized). The cells were again incubated on ice for 10 min and further centrifuged for 10 min at 3000 × g at 4 °C. The supernatant was removed, and pellets were resuspended in 5 mL cold TB buffer per 125 mL starting culture and consolidated in a single falcon tube, before adding 350 µL dimethyl sulfoxide. After another 10 min incubation on ice, 50 µL aliquots were prepared in 1.5 mL tubes and snap frozen in liquid nitrogen. Aliquots were stored at −80 °C until further use.

Chemically competent *V. natriegens* cells were transformed through the addition of plasmid DNA to an aliquot of 50 μL competent cells and incubated on ice for 30 min. After 30 min, cells were heat shocked in a water bath at 42 °C for 45 s then immediately incubated on ice for 15 min before recovery. The cells were recovered in 1 mL 37 °C warm LBv2 medium, followed by shaking at 37 °C for 1 h at 700 rpm. After recovery, the cells were pelleted by centrifugation at 3000 × g for 3 min, the supernatant was decanted, and the pellet was resuspended in the remaining ∼ 100 µL residual medium. The whole volume was plated on 37 °C warm LBv2 plates containing the appropriate antibiotic and incubated overnight at 37 °C.

### Genome engineering of *V. natriegens*

Genome engineering of *V. natriegens* was performed with the NT-CRISPR method (Stukenberg *et al*., 2022). Integration of the HiBiT tag sequence was facilitated with a gRNA that overlaps the integration site so that integration of the HiBiT tag sequence disrupts the gRNA binding site and therefore prevents CRISPR-Cas9 mediated cell killing. The plasmid pST_116 was used when a NGG PAM site was available at the integration site. The variant pST_140, carrying the almost PAM less spG Cas9 variant (Walton *et al*., 2020), was used with gRNAs relying on alternative NGN PAM sequences. The NT-CRISPR plasmids pST_116 and pST_140 were adapted for different target genes by replacing the *sfGFP* dropout with annealed oligonucleotides in a Golden Gate Assembly reaction. The annealing reactions were set up by mixing 1.5 µL of each oligonucleotide (100 µM) with 5 µL T4-DNA ligase buffer (Thermo Scientific) in a total reaction volume of 50 µL. Reactions are incubated in a heat block at 95 °C for 15 min, before the heat block was switched off to allow the samples to slowly cool down to room temperature (∼1 h). Cloning reactions with the NT-CRISPR plasmids were set up with ∼200 ng of the respective plasmid, 3 µL annealed oligonucleotides, 0.5 µL of T4-DNA Ligase (5 Weiss U/µL, Thermo Scientific) and BsaI (10 U/µL, Thermo Scientific) and 1 µL T4-DNA ligase buffer (Thermo Scientific) in a total reaction volume of 10 µL. Reactions were run in a thermocycler with 30 cycles of 37 °C (2 min) and 16 °C (5 min), followed by a final digestion step at 37 °C for 30 min and an enzyme denaturation step at 80 °C for 10 min. Oligonucleotides used for the construction of gRNAs are provided in Table S12. NT-CRISPR plasmids were first assembled in *E. coli* NEB Turbo, the plasmid DNA was then isolated and used for the transformation of *V. natriegens Δdns*.

tDNAs with homologous regions of 3 kb per arm were prepared by first constructing a tDNA template plasmid and then generating the tDNA in a PCR. The tDNA template plasmids were generated in a three fragment Gibson Assembly from two fragments comprising ∼ 3kb of sequence upstream and downstream of the integration site, amplified from genomic DNA of *V. natriegens*, as well as a backbone fragment, which was amplified from the part entry vector of the Marburg Collection pMC_V_01 (Stukenberg *et al*., 2021) or a low copy variant pMC_V_11, carrying a p15A origin of replication. The tDNA was generated in a subsequent PCR reaction (Q5 High-Fidelity DNA Polymerase, NEB) and purified using the E.Z.N.A Cycle Pure Kit (Omega Bio-Tek), according to manufacturer’s instructions. The tDNAs for the integration of the luciferase expression cassettes could not be generated with this approach, presumably due to toxicity of the homologous flanks of the integration sites or the constitutive expression of the luciferases on a multi-copy plasmid in *E. coli*. Instead, these tDNAs were created by fusion PCR. Therefore, the upstream and downstream fragments were prepared as described above. The insert fragments, carrying the luciferase expression cassette, was amplified from a level 1 plasmid which was assembled using the Marburg Collection (Stukenberg *et al*., 2021). Level 1 plasmids constructed in this study are listed in Table S6 and their plasmids maps are provided in Supplementary Data S1. For the fusion PCR to generate the final tDNA fragment, we used 50 ng of each fragment in a 25 µL PCR reaction with the outermost primers. Sequences of all primers and the templates used for the construction of tDNAs are provided in Table S13.

The NT-CRISPR protocol was performed as described before (Stukenberg *et al*., 2022). Precultures were grown overnight (16 – 17 h) at 30 °C and 200 rpm in 5 mL LBv2 with 4 µg/mL chloramphenicol and 100 µM IPTG (Roth, CAS: 367-93-1) to induce *tfoX* expression. The natural transformation was started by adding 3.5 µL of the precultures (OD_600_ ∼ 9 – 11) to 350 µL sea salt medium (28 g/L (Sigma, S9883)) with 100 µM IPTG and 100 ng of tDNA in a 1.5 mL reaction tube. Samples were briefly vortexed and then incubated statically at 30 °C for 5 h. In a subsequent step, 1 mL LBv2 without antibiotics was added to the cells. For CRISPR-Cas9 induction, 200 ng/mL ATc (Alfa Aeser, 13803-65-1) was added to the LBv2 medium. Tubes were mixed by inversion and incubated at 300 rpm and 30 °C for 1 h. Finally, 100 µL of appropriate dilutions were plated on LBv2 agar plates with 2 µg/mL chloramphenicol and 200 ng/mL ATc.

After overnight incubation at 37 °C, colonies were screened for the desired modification by colony PCR. A colony was resuspended in 20 µL of H_2_O, incubated for 10 – 15 min at 95 °C for cell lysis and cell debris was pelleted by centrifugation for 3 min at maximum speed. The PCR reaction was set up with 1 µL of the supernatant in a 12.5 µL reaction with Taq polymerase (NEB), according to manufacturer’s instruction.

Positive colonies were further verified by Sanger sequencing. Sequencing was performed by Microsynth Seqlab using PCR fragments and a primer close to the modified site.

In parallel to colony PCR screening, cells were cured from the NT-CRISPR plasmid by inoculating 5 mL of antibiotic free LBv2. After 6 – 7 h of growth at 37 °C and 200 rpm, 100 µL of a 10^-7^ dilution were plated on antibiotic free LBv2 agar plate. After overnight incubation at 37 °C, colonies were patched on LBv2 with and without 2 µg/mL chloramphenicol to check for plasmid loss. Colonies growing on the antibiotic free agar plates but not on agar plates containing chloramphenicol were considered to consist of plasmid-cured cells. Glycerol stocks of plasmid-cured strains were prepared as described above.

An overview of strains created using NT-CRISPR is shown in Table S7.

### Construction of CRISPRi plasmids

The plasmid pST_300 was used for all CRISPRi experiments with a single gRNA. pST_300 carries *dcas9* under control of the ATc inducible P_tet_ and a *gRNA* expression cassette under control of the DHBA inducible P_3b5b_. Both promoters were optimized by directed evolution for *E. coli* (Meyer *et al*., 2019). The P_3b5b_ promoter was further modified to start transcription at the first nucleotide of the *gRNA* sequence in the final plasmid configuration. The promotor sequences including information of the -35 box, -10 box, and +1 position were taken from Meyer *et al*. (2019).

The *gRNA* expression cassette carries a dropout part, which contains a *sfGFP* marker and a *sacB* expression cassette, to allow for visual identification of correctly assembled plasmids through loss of *sfGFP* and for the inhibition of cells transformed with religated pST_300 plasmid in the preparation of CRISPRi plasmids with mismatched gRNA libraries (see below). Plasmid assembly was done within the framework of the Marburg Collection (Stukenberg *et al*., 2021). pST_300 is a level 2 plasmid which was built from level 0* parts and the level 1 plasmid pST_236 using Esp3I. pST_236 was built with level 0 parts using BsaI. Golden Gate Assembly reactions were performed in a volume of 10 µL with 0.5 µL of either Esp3I (10,000 U/mL, NEB) or BsaI (10,000 U/mL Thermo Scientific), 0.5 µL T4-DNA ligase (5 Weiss U/µL, Thermo Scientific), 1 µL T4-DNA ligase buffer (Thermo Scientific) and approximately 25 fmol of each precursor plasmid. Reactions were run in a thermocycler with 30 cycles of 37 °C (5 min) and 16 °C (10 min), followed by a final digestion step at 37 °C for 60 min and an enzyme denaturation step at 80 °C for 10 min. Information about the generated level 1 and level 2 plasmids and the corresponding plasmid maps are provided in Table S6 and Supplementary Data S1.

We designed two gRNA sequences for each target gene, binding the first third of the coding sequence. The choice of gRNA sequences was done with the help of a web based tool (https://crispr-browser.pasteur.cloud/guide-rna-design) (Calvo-Villamañán *et al*., 2020). Two oligonucleotides were used for each gRNA and integrated into the CRISPRi plasmid pST_300 exactly as described above for the NT-CRISPR plasmids. Sequences of the used gRNA spacers are provided in Table S14.

### Construction of CRISPRi plasmids with mismatched gRNA libraries

For each fully-binding gRNA, we purchased three ssDNA oligonucleotide (20, 16 and 14 nucleotides), containing “N” nucleotides at positions five and ten from the PAM proximal nucleotide inside the gRNA spacer sequence. The spacer sequence of the gRNA was flanked by inward facing BsaI recognition sites, which are necessary to generate the fusion sites for the subsequent integration in pST_300. Additionally, the ssDNA oligo was flanked by primer adaptor sequences (Organick *et al*., 2018). To generate dsDNA, which is necessary for the subsequent cloning step, a 10 pM dilution of the ssDNA oligonucleotide (Table S15) was used in a Q5 PCR reaction with the primers oDS_1184 and oDS_1185, binding to the primer adaptor sequences. These primers each carry a 20-nucleotide overhang with a random sequence, to extend the final PCR fragment to ∼120 bp. Due to this extended size, the PCR fragment could be purified using the E.Z.N.A Cycle Pure Kit (Omega Bio-Tek), according to manufacturer’s instructions. The purified PCR fragments from all six oligonucleotides per target (two different gRNA sequences and three lengths) were mixed in equal amounts. This mixture was then used in threefold molar excess to 200 ng of the CRISPRi plasmid pST_300 in a 10 µL Golden Gate Assembly reaction. In this case, 0.5 µL BsaI-HFv2 (NEB, 20,000 U/mL) and 0.5 µL T4 DNA Ligase (NEB, 400,000 U/mL) were used with 1 µL T4 DNA Ligase Reaction Buffer (NEB, 10x). We found that using these enzymes resulted in higher efficiency and fidelity for the cloning of mismatched gRNA libraries. Reactions were run in a thermocycler with 30 cycles of 37 °C (5 min) and 16 °C (10 min), followed by a final digestion step at 37 °C for 60 min and an enzyme denaturation step at 80 °C for 10 min.

These cloning reactions were introduced into *E. coli* NEB Turbo through electroporation. Competent cells were prepared according to a glycerol/mannitol density step centrifugation protocol (Warren, 2011). A volume of 5 mL precultures of NEB Turbo cells was grown overnight in SOB medium (20 g/L tryptone, 5 g/L yeast extract, 0.58 g/L NaCl, 0.186 g/L KCl) at 30 °C and 200 rpm for 16-17 h and then used to inoculate 200 mL SOB in a 1 L baffled shake flask to a starting OD_600_ of 0.01. The culture was incubated and the OD_600_ carefully monitored. At an OD_600_ of approximately 0.5, the culture was distributed into four 50 mL falcon tubes and incubated for 5 min on ice. Cells were then harvested by centrifugation for 15 min at 2000 g and 4 °C in a swing rotor in a Sigma 4K15 centrifuge. All following steps were performed on ice. After centrifugation, the supernatant was first decanted and then remaining volume removed by aspiration. Each of the four pellets was gently resuspended in 10 mL of cold H_2_O. Two cell suspensions of each sample were consolidated in a clean 50 mL falcon tubes. The cold glycerol/mannitol solution (20 % glycerol (w/V), 1.5 % mannitol (g/L)) was used to create a bottom layer below the cell suspension. Approximately 12 mL of the glycerol/mannitol solution were aspirated in a 10 mL glass pipette. The glass pipette was used to pierce through the cell suspension and the glycerol/mannitol solution was slowly (∼ 20 s per tube) dispensed, resulting in an upward displacement of the cell suspension. The cells were forced through the dense glycerol/mannitol solution by another centrifugation step (2000 g, 15 min, 4 °C, acceleration and deceleration set to 2) and thereby cleaned from any remaining salts which would interfere with the electroporation. After this centrifugation step, the supernatant was carefully removed by aspiration, starting with the upper water phase, followed by the lower glycerol/mannitol phase. Each cell pellet was then resuspended in 200 µL of cold glycerol mannitol solution and cells from both falcon tubes were combined. Aliquots with 40 µL of cells were prepared in cold 1.5 mL reaction tubes and 4 µL of the Golden Gate Assembly reaction was added to the cells. The cell suspension was then transferred to a pre-chilled electroporation cuvette and after approximately 5 – 10 min electroporated in the eporator electroporator (Eppendorf) set to 1850 V. Immediately after electroporation, 1 mL of pre-warmed (37°C) SOC medium (Table S8) was added to the electroporation cuvette, mixed with the electroporated cells, and then transferred to a 1.5 mL reaction tube. Cells were recovered for 1 h at 37 °C and 700 rpm. After recovery 100 µL of the cell suspension was spread on LB agar plates (without NaCl) containing 25 µg/mL chloramphenicol and 10 % sucrose for SacB-mediated counterselection. These plates were incubated for 20 - 24 h at 30 °C to allow colonies to form. These colonies were scraped off with 2 mL of 1 % NaCl (w/v) and the resulting cell suspension was used for plasmid isolation with the E.Z.N.A. Plasmid DNA Mini Kit (Omega Bio-Tek).

Plasmid DNA resulting from this procedure was used to transform the relevant *V. natriegens* strains. After transformation, we used 48 colonies per target gene to generate glycerol stocks as described above which were then used for the respective experiments.

### Construction of CRISPRi plasmids with multiple gRNA sequences

Assembly of CRISPRi plasmids with multiple gRNA sequences was performed in two steps. In a first step, gRNA sequences were integrated into gRNA position vectors (Table S9). In the second step, the gRNA sequences from these position vectors were assembled with the CRISPRi dropout plasmid pST_301 (Fig. S15). pST_301 resembles pST_300 (used for CRISPRi with a single target) but carries an *mScarlet-I* and *sacB* expression cassette as a dropout for positions 2 – 5, according to the nomenclature of the Marburg Collection (Stukenberg *et al*., 2021).

Integration of the gRNA sequences into the position vectors was done in the same way as described before for pST_300 with the difference that kanamycin was used for selection instead of chloramphenicol. These gRNA position vectors carry a *sfGFP* and *sacB* expression cassette as a dropout to allow visual identification through loss of sfGFP production and by the inability to grow on LB sucrose due to SacB-mediated counterselection. Assembly of the final CRISPRi plasmid was done by combining the libraries of the gRNA position vectors with the pST_301 plasmid in a Golde Gate reaction as described above but with Esp3I (NEB, 10,000 U/mL) instead of BsaI. The cloning reaction was electroporated into *E. coli* NEB Turbo as described above and plated on LB sucrose agar plates with chloramphenicol to select for colonies harboring the correctly assembled plasmid. Plasmid DNA resulting from this procedure was used to transform the *V. natriegens* strain DST050 (four reporter genes integrated into the chromosome) to generate the strains for subsequent experiments.

### Measurement of growth rates and mScarlet-I fluorescence in a microplate reader

The Tecan infinite f200pro microplate reader infinite F200PRO was used for all experiments performed in 96-well plate. Plate reader protocols are provided in Table S10 and S11 for experiments without and with fluorescence measurements, respectively.

All experiments started from a precultures that were inoculated by first resuspending material from glycerol stocks in 50 µL of LBv2 (with added antibiotics if applicable) and then by adding 5 µL of this suspension to 95 µL of LBv2 (with added antibiotics if applicable). LBv2 was also used for precultures in cases where the main experiment was performed in minimal medium. The precultures was diluted 1:2000 to inoculate the 96-well plate for the experiment. The 1:2000 dilution was obtained with an intermediate dilution step (1:50) and a second dilution step (1:40). Experiments in LBv2 were performed in a total volume of 100 µL, while experiments in M9G were performed in a total volume of 150 µL to adjust to the longer experimental time and consequently for the increased evaporation over the time course of the experiment. For each well, the intermediate dilution step was performed with the exact same medium, including the exact concentrations of antibiotics and inducers as in the final experiment. For experiments using CRISPRi, precultures were induced with the same concentration of ATc and DHBA as in the main culture to induce *cas9* and *gRNA* expression and to deplete the target protein before the start of the main culture. Unless indicated otherwise, we used 200 ng/mL ATc and 100 µM DHBA as inducer concentrations for experiments in M9G and 1 µg/mL ATc and 100 µM DHBA for experiments in LBv2.

The growth rate for each culture in the 96-well plate was calculated from an exponential fit on all data points between OD_600_ values of 0.01 and 0.1, which represents the exponential growth phase.

### Measurement of mScarlet-I expression using flow cytometry

Single cell measurements of mScarlet-I signal and the effect of CRISPRi repression were performed with the BD Fortessa flow cytometer. Precultures and cultures for the measurements were grown in 96-well plates in the Tecan infinite f200pro microplate reader as described above. At an OD600 of approximately 0.1, when cells are in late exponential phase, the microplate reader measurement was aborted and cultures were diluted with PBS pH 7.4 to yield approximately 10,000 events/µL in the flow cytometry measurement. Samples were measured with the following settings: Sample flow rate = 2.0 µL/s, Sample Volume = 30 µL, Mixing Volume = 100 µL, Mixing Speed = 180 µL/s, Number of Mixes = 2, Wash Volume = 800 µL, Enable BLR = yes, BLR Period = 100. No events were excluded through gating of the raw data.

### HiBiT Assay

Abundance of the protein of interest, carrying a C-terminal fusion of the HiBiT tag, was performed with the Nano-Glo® HiBiT Lytic Detection System (Promega). Precultures and cultures for the measurements were grown in 96-well plates in the Tecan infinite f200pro microplate reader as described above. At an OD_600_ of approximately 0.1, when cells are in late exponential phase, the microplate reader measurement was aborted. Cells were transferred into a 96-well PCR plate and frozen at -80°C for at least 24 h. We found that a single freeze thaw cycle aids in cell lysis with the lysis buffer provided with the kit. Cultures were thawed immediately before the experiment at room temperature. HiBiT Assay reaction mix was freshly prepared for each experiment according to manufacturer’s instructions. All components, including thawed samples, were allowed to reach room temperature. The luminescence measurement was performed in a Tecan infinite f200pro microplate reader in black 384-well plates. For each well, 15 µL of reaction mix was combined with 15 µL of samples. To reduce spillover of luminescence, only every other row and column was used. However, as the luminescence vanishes completely after multiple days, the skipped wells could be used in later experiments. A 10 min shaking step in orbital mode with an amplitude of 3.5 mm and a frequency of 88 rpm was performed in the Tecan infinite f200pro microplate reader, before luminescence of the whole 384-well plate was performed every three minutes with an integration time of 200 ms and a settling time of 10 ms for 3 h. To correct for differences in culture density, all luminescence values were normalized by the last OD_600_ data point before the measurement was aborted. All luminescence values over the time course of 3 h were summed up to yield the final data point for the respective samples.

### Fluc and Nluc Assay

Signal of the Fluc reporter was quantified with the ONE-Glo Luciferase Assay System (Promega) and Nluc reporter was quantified with Nano-Glo Luciferase Assay System (Promega). The substrate was prepared according to manufacturer’s instructions. Samples were prepared as described for the HiBiT assay. The measurements were started by adding 15 µL of thawed samples to 15 µL of substrate in a black 384-well plate. Measurement and data analysis was done as described for the HiBiT assay.

### Rluc Assay

Signal of the Fluc reporter was quantified with the Renilla Luciferase Assay System (Promega). The substrate was prepared according to manufacturer’s instructions. Samples were prepared as described for the HiBiT assay. After thawing, the samples were diluted with 2 x Luciferase Assay Lysis Buffer (diluted from 5 x with H_2_O) and incubated for at least 10 min at room temperature. Due to the shorter signal half-life time compared to the other luciferase systems, the assay was performed differently than for the other luciferases. 15 µL of the samples (after dilution with lysis buffer) were transferred to wells of a black 384-well plate. 15 µL of the substrate was added to the cells with 10 s delay between wells. A single luminescence measurement was done also with a 10 s delay between wells, to ensure the same time delay from addition of substrate to measurement for each well. Six samples per run were processed to minimize loss of signal before the measurements.

### Calculation of probability distribution

A probability distribution was generated to predict the expectable number of unique variants from gRNA libraries. Therefore, a custom MATLAB script was created with the following steps. First, a random number between 1 and 96 was generated, each representing one of the 96 theoretically possible variants from the gRNA library. This was repeated 48 times, representing the 48 sequenced variants. Next, the number of unique variants was determined. Finally, this was repeated 10,000 times to generate the histogram for the probability of obtaining a specific number of unique variants.

## Supporting Information

Figure S1: Effect of inducer concentrations on repression strength in LBv2.

Figure S2: Effect of inducer concentrations on growth rate in LBv2.

Figure S3: Inducibility of CRISPRi system and comparison with control constructs in LBv2.

Figure S4: Effect of inducer concentrations on mScarlet-I signal with non-binding control gRNA.

Figure S5: Analyzing the mScarlet-I signal resulting from different combinations of DHBA and ATc at the single cell level in LBv2.

Figure S6: Repression strengths with gRNAs featuring mismatches and different spacer lengths in LBv2.

Figure S7: Probability distribution for number of unique variants.

Figure S8: Comparing single-cell distribution from mismatched and truncated gRNAs with limiting inducer concentrations.

Figure S9: Correlation between Fluc and HiBiT signal.

Figure S10: Testing the effect of additional gRNAs on growth rate at different ATc concentrations.

Figure S11: Crosstalk between reporter signals.

Figure S12: Correlation between reporter signals from a multi gRNA library experiment (Fig. 3c).

Figure S13: Growth curves of HiBiT fusion strains.

Figure S14: Growth curves of *argA* and *metE* deletion strains.

Table S1: Sequencing results of mScarlet-I gRNA library variants.

Table S2: Recipe for buffered M9 minimal medium with 0.4 % (w/w) glucose (M9G).

Table S3: Recipe 5X M9 salts solution.

Table S4: Composition of 100X trace elements solution.

Table S5: Preparation of 100x trace elements solution.

Table S6: Level 1 and level 2 plasmids assembled in this study.

Table S7: *V. natriegens* strains with chromosomal modification

Table S8: Recipe for SOC medium

Table S9: gRNA position vectors for assembly of multi gRNA CRISPRi plasmids.

Table S10: Protocol for microplate reader measurements without fluorescence measurement

Table S11: Protocol for microplate reader measurements with fluorescence measurement

Table S12: Oligonucleotides used to assemble gRNA sequences for NT-CRISPR

Table S13: Oligonucleotides used for the construction of tDNA template plasmids and tDNAs

Table S14: Oligonucleotides used to assemble gRNA sequences for CRISPRi

Table S15: ssDNA templates for the generation of gRNA libraries Data S1: Plasmid maps

## Author Information

### Corresponding author

**Anke Becker** - *Center for Synthetic Microbiology, Philipps-Universität Marburg, Marburg, Germany and Department of Biology, Philipps-Universität Marburg, Marburg, Germany*

### Authors

**Daniel Stukenberg** - Center for Synthetic Microbiology, Philipps-Universität Marburg, Marburg, Germany and Department of Biology, Philipps-Universität Marburg, Marburg, Germany

**Anna Faber** - Center for Synthetic Microbiology, Philipps-Universität Marburg, Marburg, Germany and Department of Biology, Philipps-Universität Marburg, Marburg, Germany Current address: School of Molecular Sciences, The University of Western Australia, Crawley, Australia

### Author contributions

D.S., A.F., and A.B. conceived this project. A.F. performed proof-of-concept experiments and D.S. performed all experiments described in this publication. D.S. constructed all strains and plasmids used in this study. D.S. analyzed the data and created all figures. D.S., A.F., and A.B. wrote the manuscript. A.B. supervised the study.

## Supporting information

Plasmid maps

## Acknowledgements

This work was funded by the State of Hesse (Germany) through the LOEWE cluster Diffusible signals. D.S. received funding through the International Max Planck Research School for Environmental, Cellular, and Molecular Microbiology (IMPRS-Mic). We thank Prof. Christopher A. Voigt for the Marionette Sensor Collection, which was supplied through Addgene (Addgene Kit #1000000137) and Silvia González Sierra for her assistance with flow cytometry measurements. Furthermore, we thank Reza Rohani and Lucas Coppens for sharing their insights into the metabolism of *V. natriegens*.

## References

Calvo-Villamañán, A. et al. (2020) ‘On-target activity predictions enable improved CRISPR–dCas9 screens in bacteria’, Nucleic acids research, 48(11), pp. e64–e64. Available at: 10.1093/nar/gkaa294.

Ceroni, F. et al. (2018) ‘Burden-driven feedback control of gene expression’, Nature Methods, 15(5), pp. 387–393. Available at: 10.1038/nmeth.4635.

Cho, S. et al. (2018) ‘High-Level dCas9 Expression Induces Abnormal Cell Morphology in *Escherichia coli*’, ACS Synthetic Biology, 7(4), pp. 1085–1094. Available at: 10.1021/acssynbio.7b00462.

Coppens, L. et al. (2023) ‘*Vibrio natriegens* genome-scale modeling reveals insights into halophilic adaptations and resource allocation’, Molecular Systems Biology, 19(4), p. e10523. Available at: 10.15252/msb.202110523.

Dalia, T.N. et al. (2017) ‘Multiplex Genome Editing by Natural Transformation (MuGENT) for Synthetic Biology in Vibrio natriegens’, ACS Synthetic Biology, 6(9), pp. 1650–1655. Available at: 10.1021/acssynbio.7b00116.

Eagon, R.G. (1961) ‘Generation Time of Less Than 10 Minutes’, pp. 1961–1962.

Fontana, J. et al. (2018) ‘Regulated Expression of sgRNAs Tunes CRISPRi in *E. coli*’, Biotechnology Journal, 13(9), p. 1800069. Available at: 10.1002/biot.201800069.

Fu, Y. et al. (2014) ‘Improving CRISPR-Cas nuclease specificity using truncated guide RNAs’, Nature biotechnology, 32(3), pp. 279–284. Available at: 10.1038/nbt.2808.

Hawkins, J.S. et al. (2020) ‘Mismatch-CRISPRi Reveals the Co-varying Expression-Fitness Relationships of Essential Genes in Escherichia coli and Bacillus subtilis’, Cell Systems, 11(5), pp. 523–535.e9. Available at: 10.1016/j.cels.2020.09.009.

Hoff, J. et al. (2020) ‘*Vibrio natriegens*: an ultrafast-growing marine bacterium as emerging synthetic biology chassis’, Environmental microbiology, 22(10), pp. 4394–4408. Available at: 10.1111/1462-2920.15128.

Hoffart, E. et al. (2017) ‘High substrate uptake rates empower *Vibrio natriegens* as production host for industrial biotechnology’, Applied and environmental microbiology, 83(22), pp. 1–10. Available at: 10.1128/AEM.01614-17.

Kim, S.K. et al. (2017) ‘CRISPR interference-guided multiplex repression of endogenous competing pathway genes for redirecting metabolic flux in *Escherichia coli*’, Microbial Cell Factories, 16(1), p. 188. Available at: 10.1186/s12934-017-0802-x.

Lee, H.H. et al. (2019) ‘Functional genomics of the rapidly replicating bacterium *Vibrio natriegens* by CRISPRi’, Nature Microbiology, 4(7), pp. 1105–1113. Available at: 10.1038/s41564-019-0423-8.

Long, C.P. et al. (2017) ‘Metabolism of the fast-growing bacterium *Vibrio natriegens* elucidated by (13)C metabolic flux analysis’, Metabolic engineering, 44, pp. 191–197. Available at: 10.1016/j.ymben.2017.10.008.

Meyer, A.J. et al. (2019) ‘*Escherichia coli* "Marionette" strains with 12 highly optimized small-molecule sensors’, Nature Chemical Biology, 15(2), pp. 196–204. Available at: 10.1038/s41589-018-0168-3.

O’Brien, E.J., Utrilla, J. and Palsson, B.O. (2016) ‘Quantification and Classification of *E. coli* Proteome Utilization and Unused Protein Costs across Environments’, PLOS Computational Biology, 12(6), p. e1004998. Available at: 10.1371/journal.pcbi.1004998.

Organick, L. et al. (2018) ‘Random access in large-scale DNA data storage’, Nature biotechnology, 36(3), pp. 242–248. Available at: 10.1038/nbt.4079.

Qi, L.S. et al. (2013) ‘Repurposing CRISPR as an RNA-Guided Platform for Sequence-Specific Control of Gene Expression’, Cell, 152(5), pp. 1173–1183. Available at: 10.1016/j.cell.2013.02.022.

Sander, T. et al. (2019) ‘Allosteric Feedback Inhibition Enables Robust Amino Acid Biosynthesis in *E. coli* by Enforcing Enzyme Overabundance’, Cell Systems, 8(1), pp. 66–75.e8. Available at: 10.1016/j.cels.2018.12.005.

Schwinn, M.K. et al. (2018) ‘CRISPR-Mediated Tagging of Endogenous Proteins with a Luminescent Peptide’, ACS Chemical Biology, 13(2), pp. 467–474. Available at: 10.1021/acschembio.7b00549.

Stukenberg, D. et al. (2021) ‘The Marburg Collection: A Golden Gate DNA Assembly Framework for Synthetic Biology Applications in *Vibrio natriegens*’, ACS Synthetic Biology, 10(8), pp. 1904–1919. Available at: 10.1021/acssynbio.1c00126.

Stukenberg, D. et al. (2022) ‘NT-CRISPR, combining natural transformation and CRISPR-Cas9 counterselection for markerless and scarless genome editing in *Vibrio natriegens*’, Communications Biology, 5(1), p. 265. Available at: 10.1038/s42003-022-03150-0.

Thoma, F. et al. (2022) ‘Metabolic engineering of *Vibrio natriegens* for anaerobic succinate production’,

*Microbial* *Biotechnology*, 15(6), pp. 1671–1684. Available at: 10.1111/1751-7915.13983.

Thoma, F. and Blombach, B. (2021) ‘Metabolic engineering of Vibrio natriegens’, Essays in Biochemistry. Edited by D. Mattanovich and P. Ivan Nikel, 65(2), pp. 381–392. Available at: 10.1042/EBC20200135.

Tian, T. et al. (2019) ‘Redirecting Metabolic Flux via Combinatorial Multiplex CRISPRi-Mediated Repression for Isopentenol Production in *Escherichia coli*’, ACS Synthetic Biology, 8(2), pp. 391–402. Available at: 10.1021/acssynbio.8b00429.

Tietze, L. et al. (2022) ‘Identification and Cross-Characterisation of Artificial Promoters and 5′ Untranslated Regions in Vibrio natriegens’, Frontiers in Bioengineering and Biotechnology, 10. Available at: https://www.frontiersin.org/articles/10.3389/fbioe.2022.826142 (Accessed: 13 September 2023).

Tschirhart, T. et al. (2019) ‘Synthetic Biology Tools for the Fast-Growing Marine Bacterium *Vibrio natriegens*’, ACS Synthetic Biology, 8(9), pp. 2069–2079. Available at: 10.1021/acssynbio.9b00176.

Vigouroux, A. et al. (2018) ‘Tuning dCas9’s ability to block transcription enables robust, noiseless knockdown of bacterial genes’, Molecular systems biology, 14(3), pp. e7899–e7899. Available at: 10.15252/msb.20177899.

Walton, R.T. et al. (2020) ‘Unconstrained genome targeting with near-PAMless engineered CRISPR-Cas9 variants’, Science, 368(6488), p. 290 LP – 296. Available at: 10.1126/science.aba8853.

Wang, T. et al. (2018) ‘Pooled CRISPR interference screening enables genome-scale functional genomics study in bacteria with superior performance’, Nature Communications, 9(1), p. 2475. Available at: 10.1038/s41467-018-04899-x.

Wang, Z. et al. (2020) ‘Melanin produced by the fast-growing marine bacterium *Vibrio natriegens* through heterologous biosynthesis: Characterization and application’, Applied and environmental microbiology, 86(5), pp. e02749–19. Available at: 10.1128/AEM.02749-19.

Warren, D.J. (2011) ‘Preparation of highly efficient electrocompetent *Escherichia coli* using glycerol/mannitol density step centrifugation’, Analytical Biochemistry, 413(2), pp. 206–207. Available at: 10.1016/j.ab.2011.02.036.

Weinstock, M.T. et al. (2016) ‘*Vibrio natriegens* as a fast-growing host for molecular biology’, Nature Methods, 13(10), pp. 849–851. Available at: 10.1038/nmeth.3970.

Xu, L. et al. (2006) ‘Average Gene Length Is Highly Conserved in Prokaryotes and Eukaryotes and Diverges Only Between the Two Kingdoms’, Molecular Biology and Evolution, 23(6), pp. 1107–1108. Available at: 10.1093/molbev/msk019.

Xu, Y. et al. (2000) ‘Evolution of Arginine Biosynthesis in the Bacterial Domain: Novel Gene-Enzyme Relationships from Psychrophilic Moritella Strains (*Vibrionaceae*) and Evolutionary Significance of N-α-Acetyl Ornithinase’, Journal of Bacteriology, 182(6), pp. 1609–1615. Available at: 10.1128/jb.182.6.1609-1615.2000.

