## Supplementary Material for "graded-CRISPRi, a novel tool for tuning the strengths of CRISPRi-mediated knockdowns in *Vibrio natriegens* using gRNA libraries"

### Contents:

Figure S1: Effect of inducer concentrations on repression strength in LBv2

Figure S2:  Effect of inducer concentrations on growth rate in LBv2

Figure S3: Inducibility of CRISPRi system and comparison with control constructs in LBv2

Figure S4: Effect of inducer concentrations on mScarlet-I signal with non-binding control gRNA

Figure S5: Analyzing the mScarlet-I signal resulting from different combinations of DHBA and ATc at the single cell level in LBv2

Figure S6: Repression strengths with gRNAs featuring mismatches and different spacer lengths in LBv2

Figure S7: Probability distribution for number of unique variants

Figure S8: Comparing single-cell distribution from mismatched and truncated gRNAs with limiting inducer concentrations

Figure S9: Correlation between Fluc and HiBiT signal

Figure S10: Testing the effect of additional gRNAs on growth rate at different ATc concentrations

Figure S11: Crosstalk between reporter signals

Figure S12: Correlation between reporter signals from a multi gRNA library experiment (Fig. 3c)

Figure S13: Growth curves of HiBiT fusion strains

Figure S14: Growth curves of *argA* and *metE* deletion strains

Table S1: Sequencing results of mScarlet-I gRNA library variants

Table S2: Recipe for buffered M9 minimal medium with 0.4 % (w/w) glucose (M9G)

Table S3: Recipe 5X M9 salts solution

Table S4: Composition of 100X trace elements solution

Table S5: Preparation of 100x trace elements solution

Table S6: Level 1 and level 2 plasmids assembled in this study

Table S7: *V. natriegens* strains with chromosomal modification

Table S8: Recipe for SOC medium

Table S9: gRNA position vectors for assembly of multi gRNA CRISPRi plasmids

Table S10: Protocol for microplate reader measurements without fluorescence measurement

Table S11: Protocol for microplate reader measurements with fluorescence measurement

### Supplementary Figures

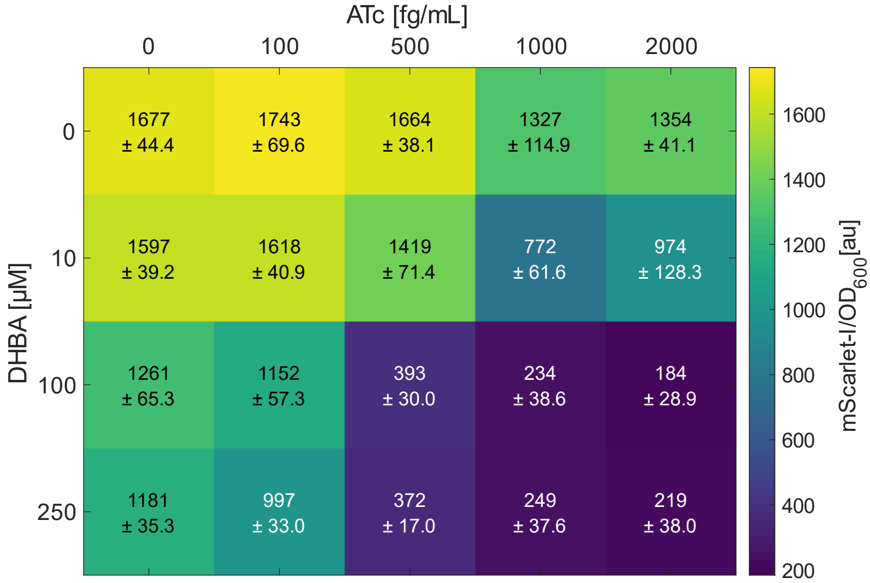


**Figure S1: Effect of inducer concentrations on repression strength in LBv2.** Colors in the heatmap indicate mScarlet-I fluorescence signal. Embedded text reports the mean mScarlet-I signal, as well as the standard deviation from the mean. This data is based on two independent experiments with three biological replicates. Experiments were performed in LBv2 with CRISPRi plasmid pST_300 and strain DST018.

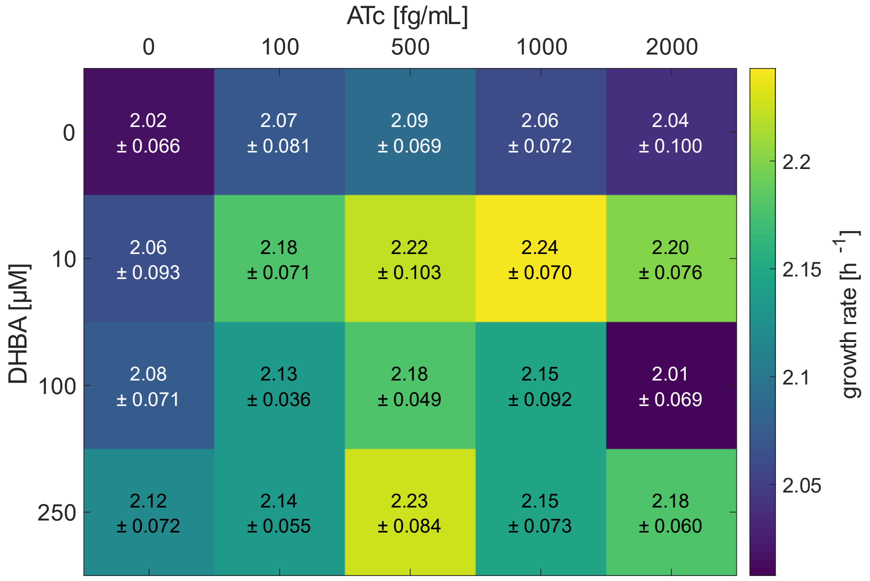


**Figure S2: Effect of inducer concentrations on growth rate in LBv2.** Colors in the heatmap indicate the growth rate of the cultures of strain DST018 harboring the CRISPRi plasmid pST_300 in LBv2. Embedded text reports the mean growth rate, as well as the standard deviation from the mean. This data is based on two independent experiments with three biological replicates.

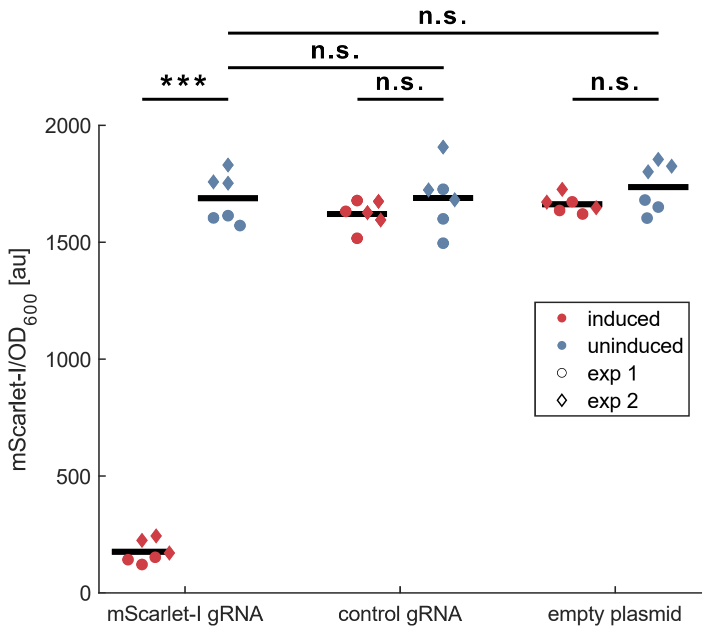


**Figure S3: Inducibility of CRISPRi system and comparison with control constructs in LBv2.** Induced samples (1000 fg/mL ATc, 100 µM DHBA) and uninduced samples are shown in red and blue, respectively. Data points from the first independent experiment are shown as circles and data from the second experiment are displayed as diamonds. Control gRNA refers to a gRNA targeting *dns*, which is deleted in this strain. The experiment was performed in LBv2 with the CRISPRi plasmid pST_300 in strain DST018. Significances were calculated with a two-sample t-test. n.s.: $p > 0.05$, *: $p > 0.01$, **: $p > 0.001$, ***: $p < 0.001$.

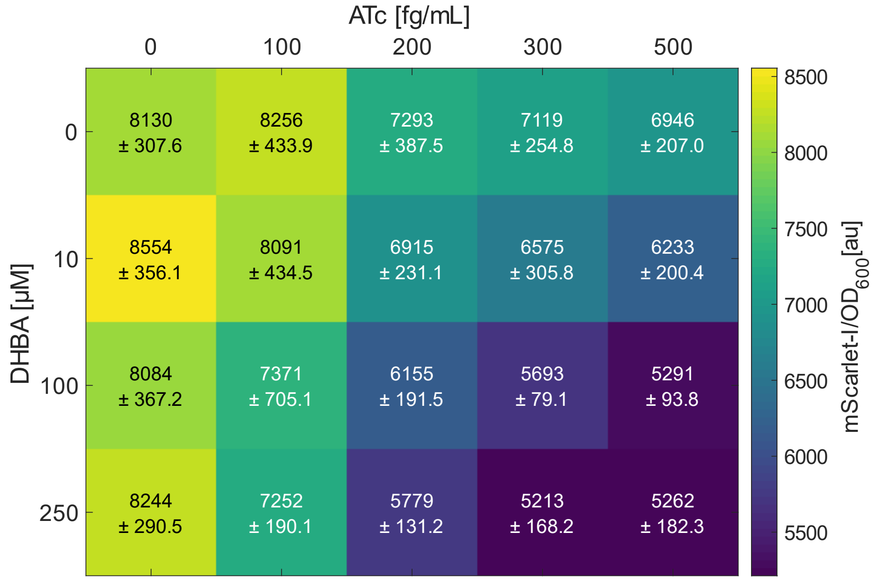


**Figure S4: Effect of inducer concentrations on mScarlet-I signal with non-binding control gRNA.** Colors in the heatmap indicate mScarlet-I fluorescence signal. Embedded text represents the mean mScarlet-I signal, as well as the standard deviation from the mean. This data is based on two independent experiments with three biological replicates. Experiments were performed in M9G with the CRISPRi plasmid pST_300 and strain DST018. A gRNA targeting *dns* (deleted in this strain) was used.

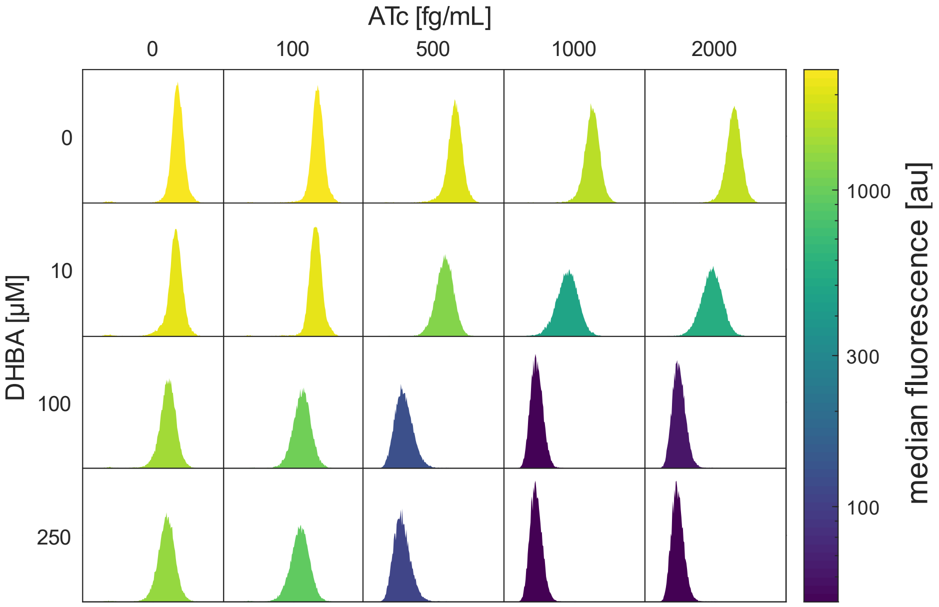


**Figure S5: Analyzing the mScarlet-I signal resulting from different combinations of DHBA and ATc at the single cell level in LBv2.** Fluorescence was measured with a flow cytometer. Displayed is a representative histogram from six measurements (three biological replicates, and two independent experiments). The color of the histograms represents the median fluorescence. Samples were drawn from cultures grown in LBv2 with the CRISPRi plasmid pST_300 in the strain DST018.

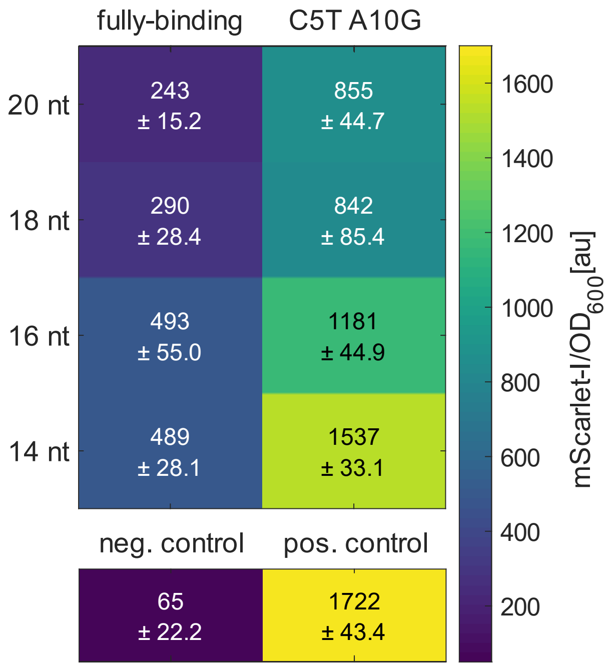


**Figure S6: Repression strengths with gRNAs featuring mismatches and different spacer lengths in LBv2.** C5T and A10G indicate mismatches to the target sequence and are counted form the PAM proximal nucleotide of the spacer sequence. - strain: strain without integrated mScarlet-I cassette (DST016). + strain: strain integrated mScarlet-I cassette (DST018) with a CRISPRi plasmid with a non-binding control gRNA. Colors in the heatmap indicate mScarlet-I fluorescence signal. Embedded text represents the mean mScarlet-I signal, as well as the standard deviation from the mean. This data is based on two independent experiments with three biological replicates. Experiments were performed in LBv2 with the CRISPRi plasmid pST_300.

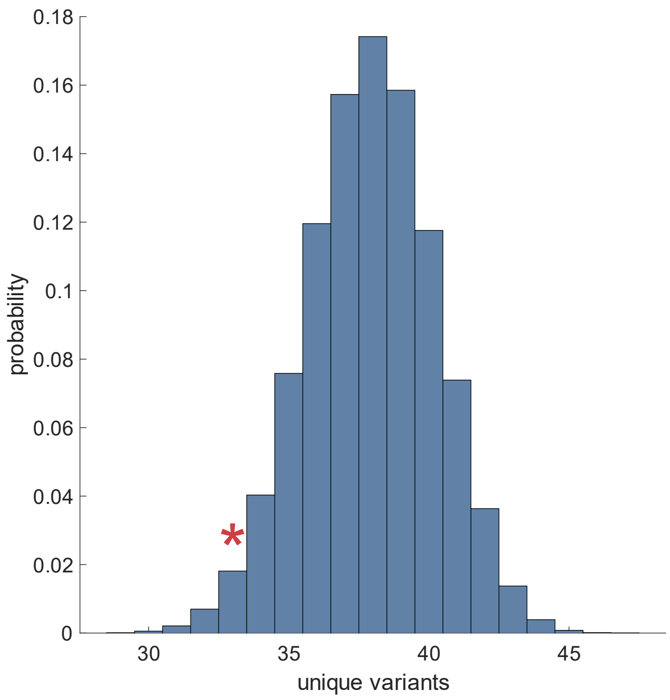


**Figure S7: Probability distribution for number of unique variants**. Histogram depicting the probability distribution of obtaining a specific number of unique variants when 48 clones are tested from 96 theoretically possible unique variants. The red asterisk indicates the number of unique variants obtained when testing 48 colonies from the gRNA library targeting mScarlet-I.

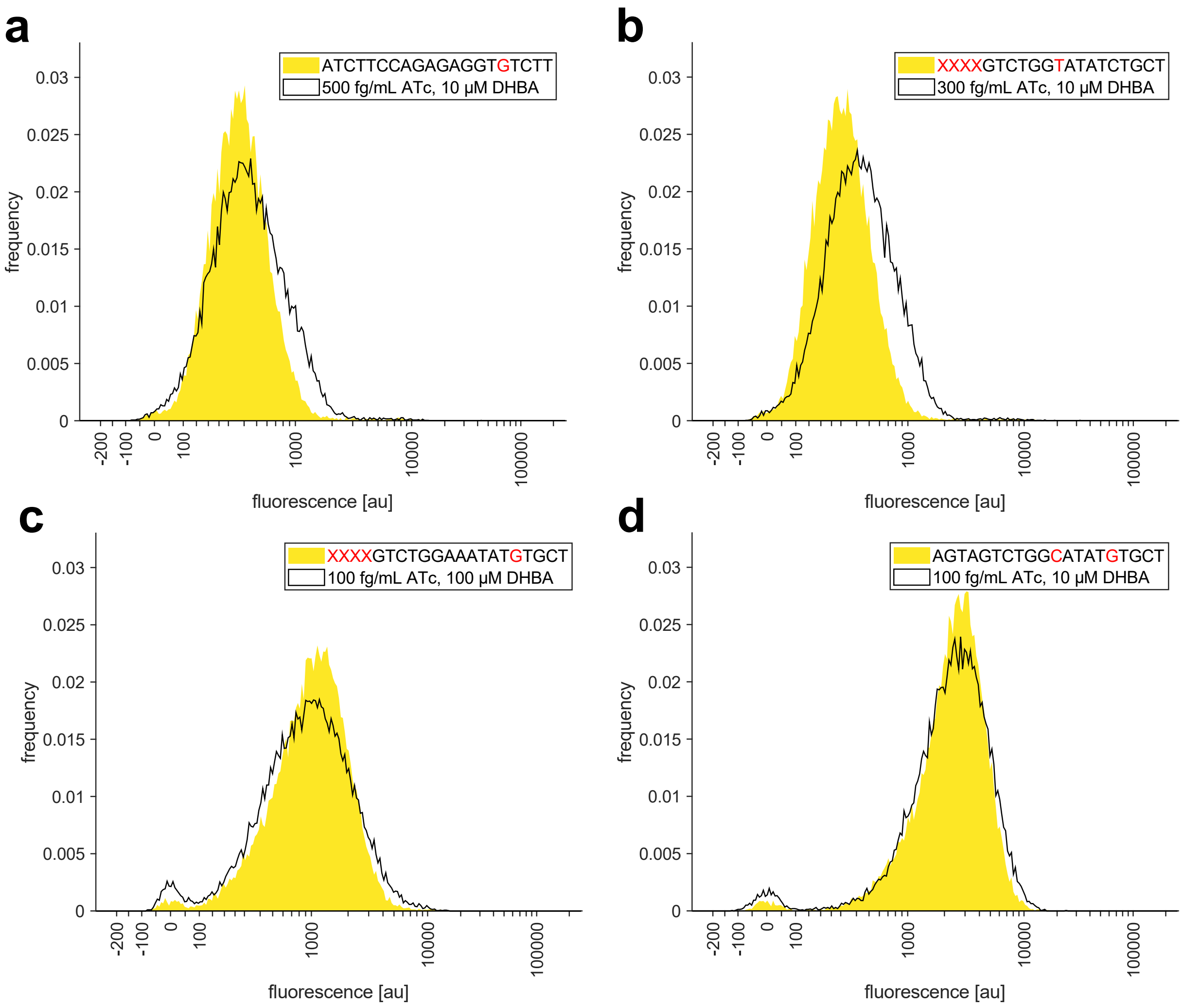


**Figure S8: Comparing single-cell distribution from mismatched and truncated gRNAs with limiting inducer concentrations.** Filled yellow histograms represent result from fully induced CRISPRi system (200 fg/mL ATc, 100 µM DHBA) with mismatched and truncated gRNAs. Red letters in the legend indicate mismatches in the gRNA variants. Red "X" letters indicate truncations. Black line represents the outline of a histogram from samples with fully-binding gRNA against mScarlet-I but with limiting inducer concentrations (specified in the legend). Experiments were performed in M9G with the CRISPRi plasmid pST_300 and strain DST018.

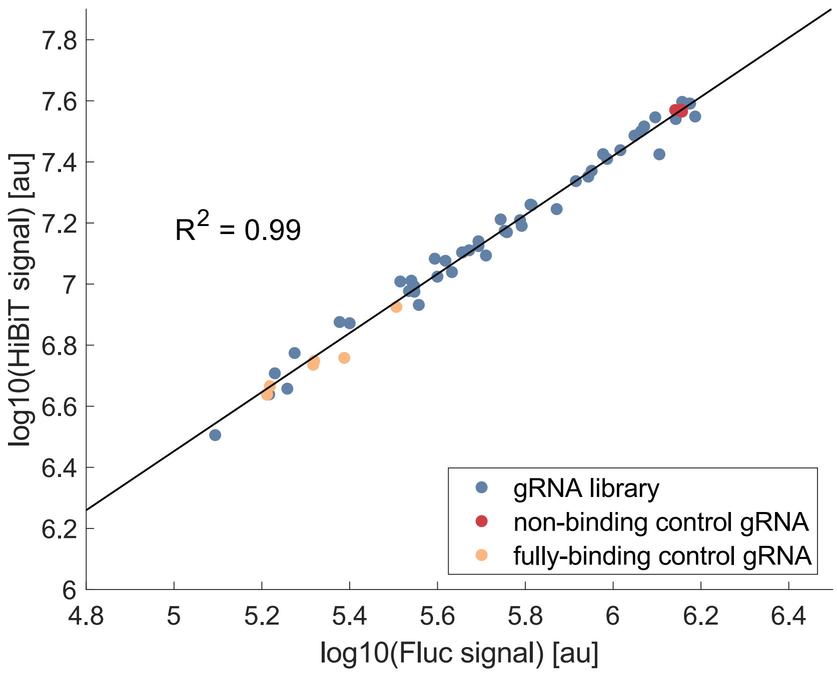


**Figure S9: Correlation between Fluc and HiBiT signal.** The experiment is based on a gRNA library targeting Fluc in a strain with a chromosomally integrated Fluc-HiBiT expression cassette (DST046). Each data point represents a single measurement from a variant of the gRNA library (blue), the non-binding control gRNA (red) or the fully-binding gRNA (yellow). $R^2$ is derived from a linear regression of the log10 of the luminescence values from Fluc and HiBiT. Experiments performed in M9G with induction of the CRISPRi system (200 fg/mL ATc, 100 µM DHBA).

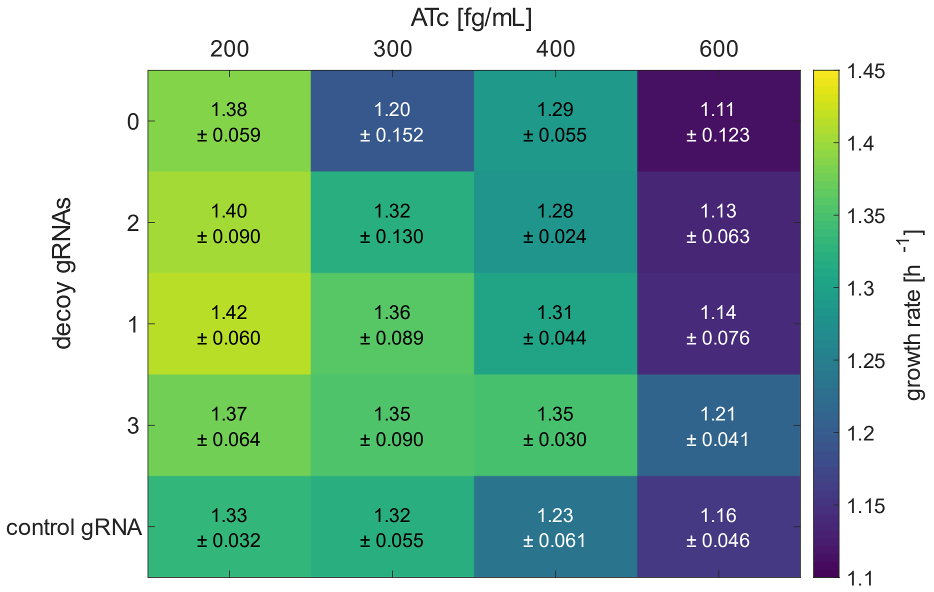


**Figure S10: Testing the effect of additional gRNAs on growth rate at different ATc concentrations.** CRISPRi plasmids encoding a gRNA targeting mScarlet-I and up to three decoy gRNAs (targeting *dns* = non-binding gRNA) were tested at different ATc concentrations and the same DHBA concentration (100 µM). Colors in the heatmap indicate the growth rate of the cultures. Embedded text represents the mean growth rate, as well as the standard deviation from the mean. This data is based on two independent experiments with three biological replicates. Experiments were performed in M9G in strain DST018.

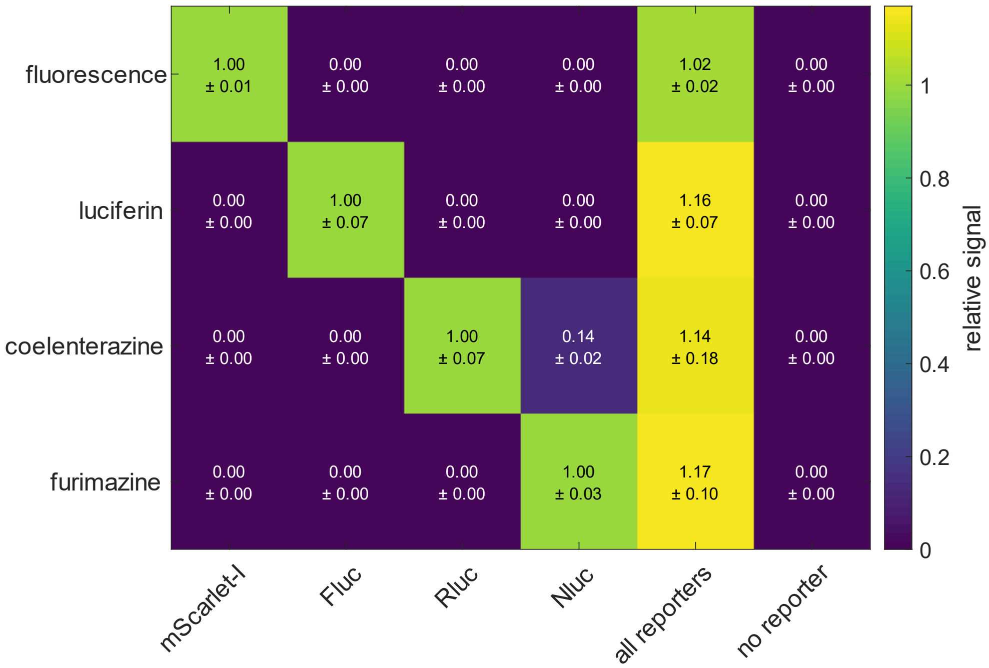


**Figure S11: Crosstalk between reporter signals.** Strains encoding one of the reporter proteins mScarlet-I, Fluc, Rluc, Nluc, a strain with all four reporter genes and a strain without any reporter gene were tested. The heatmap shows the mean signal from three biological replicates, relative to the respective single reporter strain, from either fluorescence or the respective luciferase substrates (see methods chapter). Embedded text reports the mean relative signal, as well as the standard deviation from the mean. This data is derived from three biological replicates and the samples were obtained from cultures grown in M9G.

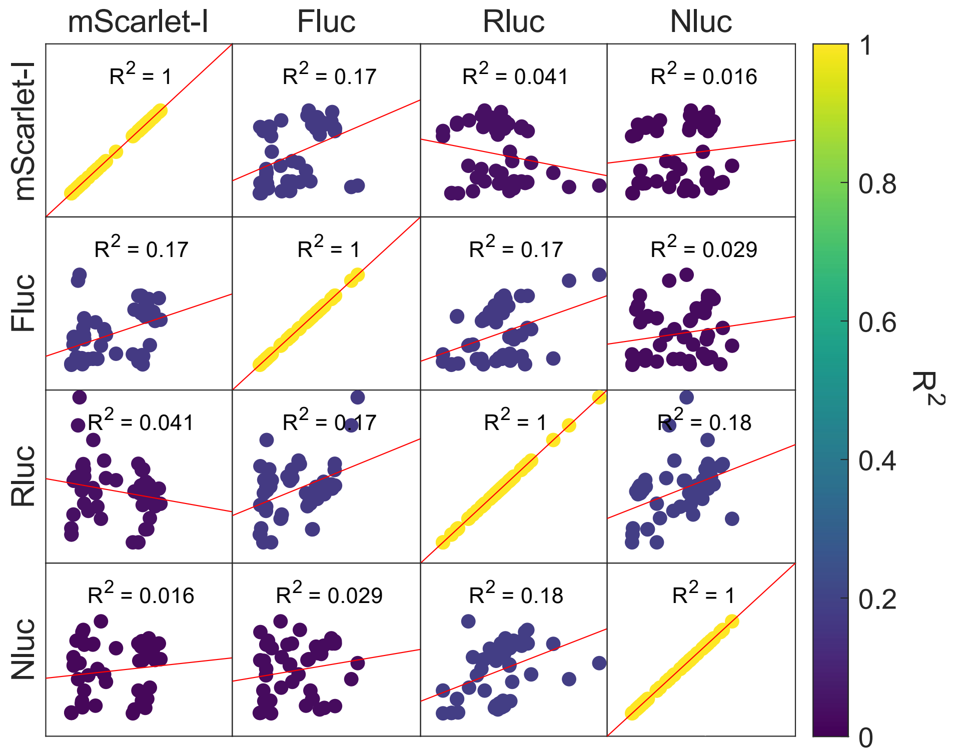


**Figure S12: Correlation between reporter signals from a multi gRNA library experiment (Fig. 3c).** Relative signals from the respective reporter genes are shown as a scatter plot. Dots are colored according to the $R^2$ value obtained from a linear regression between the relative signals of both reporters.

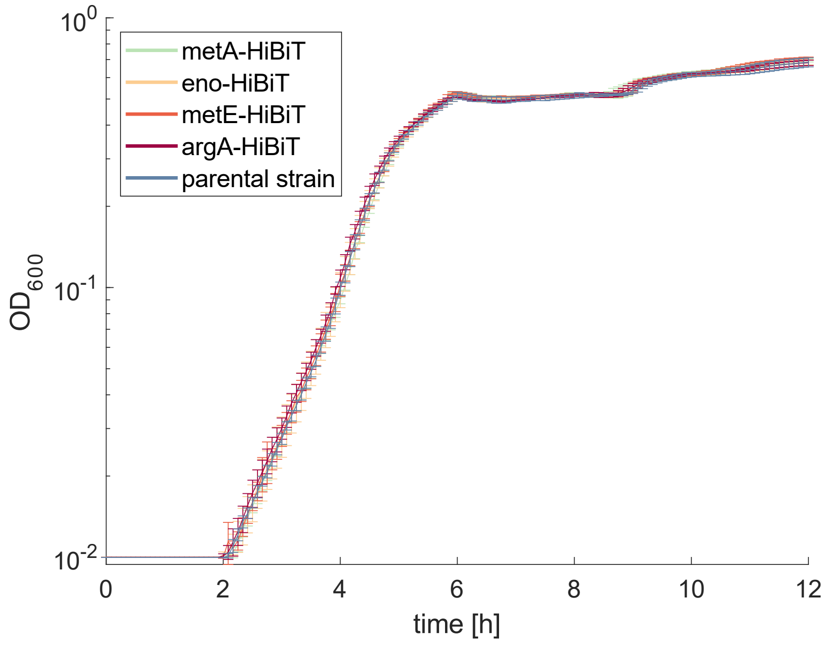


**Figure S13: Growth curves of HiBiT fusion strains.** Growth of strains with fusion of HiBiT sequence to native genes in comparison to the parental strain (DST016). The lines reflect the mean from three biological replicates and error bars indicate standard deviation from the mean. The experiment was performed in M9G.

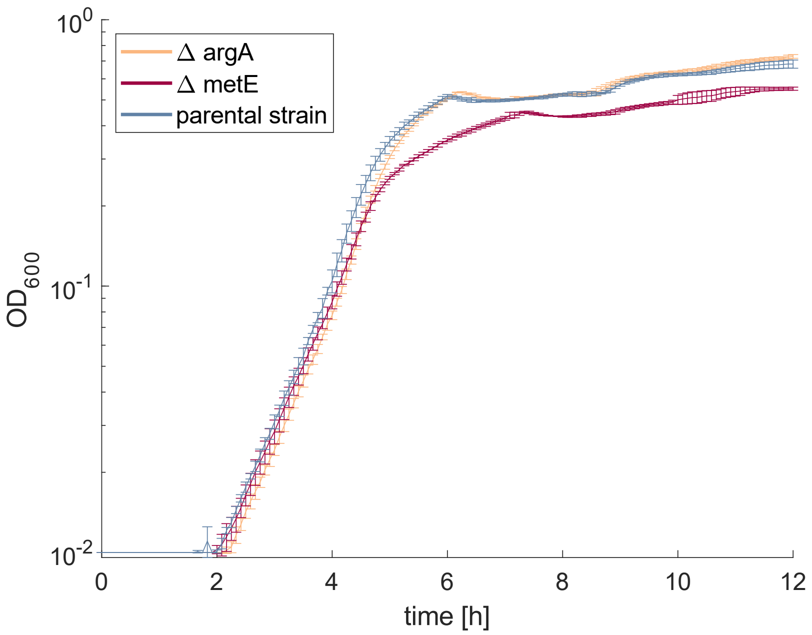


**Figure S14: Growth curves of *argA* and *metE* deletion strains.** Growth of strains deletion of *argA* and *metE* in comparison to the parental strain (DST016). The lines reflect the mean from three biological replicates and error bars indicate the standard deviation from the mean. The experiment was performed in M9G.

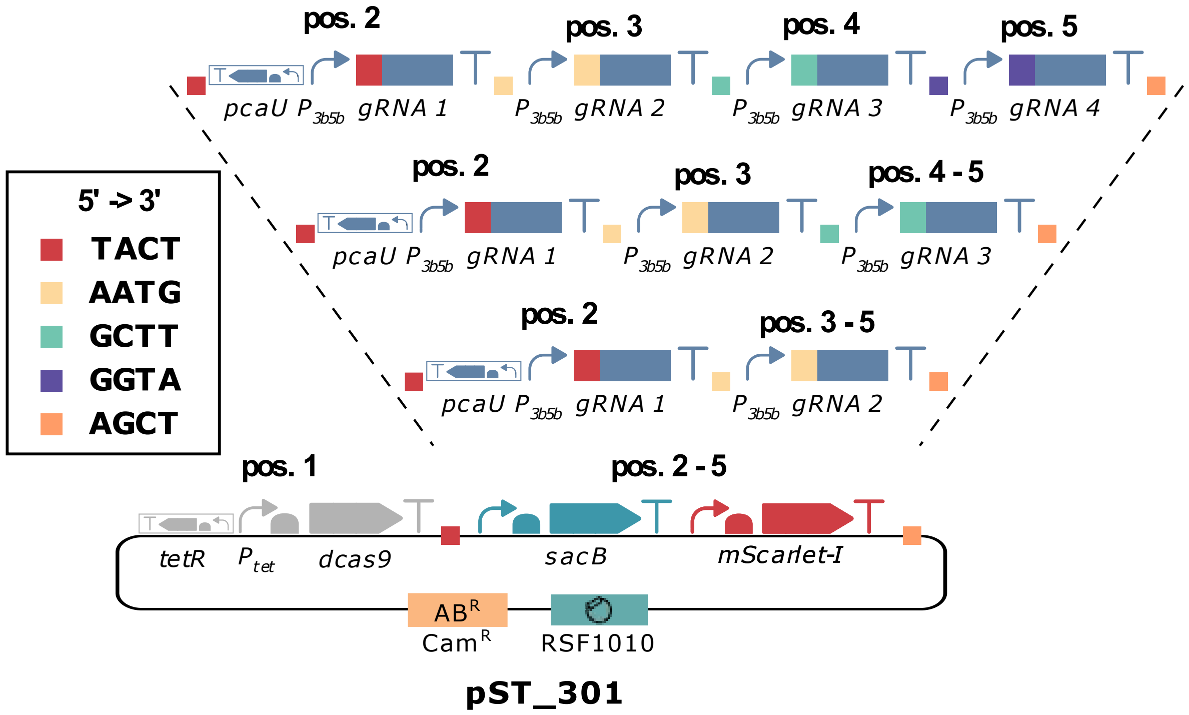


**Figure S15: Assembly scheme for CRISPRi plasmids with multiple gRNAs.** pST_301 carries a *sacB-mScarlet-I* dropout part, which is replaced with the respective gRNA expression cassettes. Different gRNA position vectors are used, depending on the number of desired gRNAs in the final construct.

### Supplementary Tables

**Table S1: Sequencing results of mScarlet-I gRNA library variants.**

| Clone | gRNA | length | mismatch |
|---|---|---|---|
| 1 | 1 | 14 | A10C |
| 2 | 1 | 16 | C5G |
| 3 | 1 | 20 | C5G, A10C |
| 4 | 2 | 20 | A5G |
| 5 | 1 | 16 | C5A, A10G |
| 6 | 1 | 14 | C5G, A10T |
| 7 | 1 | 16 | C5A, A10G |
| 8 | 1 | 20 | C5A, A10G |
| 9 | 1 | 20 | A10C |
| 10 | 2 | 20 | A5T, G10T |
| 11 | 2 | 20 | none |
| 12 | 2 | 20 | A5G, G10T |
| 13 | 1 | 16 | A5G, A10G |
| 14 | 1 | 20 | C5G, A10C |
| 15 | 2 | 20 | A5T, G10A |
| 16 | 1 | 14 | C5T, A10G |
| 17 | 2 | 16 | A5C, G10T |
| 18 | 1 | 20 | none |
| 19 | 2 | 20 | A5T, G10C |
| 20 | 1 | 14 | A10C |
| 21 | 1 | 16 | C5G, A10C |
| 22 | 1 | 16 | C5T, A10G |
| 23 | 2 | 20 | A5T, G10T |
| 24 | 1 | 16 | A10C |
| 25 | 1 | 16 | A10T |
| 26 | 2 | 20 | G10A |
| 27 | 1 | 20 | C5T |
| 28 | 1 | 14 | A10G |
| 29 | 2 | 20 | A5T, G10A |
| 30 | 2 | 14 | A5T, G10T |
| 31 | 2 | 14 | A5T |
| 32 | 1 | 16 | A10C |
| 33 | 2 | 16 | A5C |
| 34 | 2 | 20 | A5G, G10A |
| 35 | 1 | 16 | A10C |
| 36 | 2 | 16 | A5T |
| 37 | 2 | 14 | A5T |
| 39 | 2 | 20 | A5T, G10A |
| 40 | 1 | 14 | C5G, A10G |
| 41 | 1 | 16 | C5T, A10G |
| 43 | 1 | 20 | C5A, A10G |
| 44 | 1 | 20 | none |
| 45 | 2 | 20 | A5G, G10A |

| | | | |
|---|---|---|---|
| 46 | 1 | 20 | A10T |
| 47 | 1 | 14 | A10G |
| 48 | 1 | 20 | none |
| 49 | 2 | 16 | A5C, G10C |
| 50 | 2 | 14 | A5G, G10T |

**Table S2: Recipe for buffered M9 minimal medium with 0.4 % (w/w) glucose (M9G).**

| Component | Volume |
|---|---|
| NaCl solution (20% w/w) | 5 mL |
| 5X M9 salts solution | 10 mL |
| Glucose solution (20% w/w) | 1 mL |
| 100X trace elements solution | 500 µL |
| $H_2O$ | Add up to 50 mL |
| $MgSO_4$ (1 M) | 50 µL |
| $CaCl_2$ (1M) | 15 µL |
| $K_2HPO_4$ (1 M) | 2.5 mL |
| $KH_2PO_4$ (1 M) | 2.5 mL |

**Table S3: Recipe 5X M9 salts solution.**

| Component | Concentration |
|---|---|
| $KH_2PO_4$ | 15 g/L |
| $Na_2HPO_4 + 7\ H_2O$ | 64 g/L |
| NaCl | 2.5 g/L |
| $NH_4Cl$ | 5 g/L |

**Table S4: Composition of 100X trace elements solution.**

| Component | Concentration |
|---|---|
| EDTA | 5 g /L |
| $FeCl_3 + 6H_2O$ | 830 mg/L |
| $ZnCl_2$ | 84 mg/L |
| $CuCl_2 + 2H_2O$ | 13 mg/L |
| $CoCl_2 + 2H_2O$ | 10 mg/L |
| $H_3BO_3$ | 10 mg/L |
| $MnCl_2 + 4H_2O$ | 1.6 mg/L |

**Table S5: Preparation of 100x trace elements solution.** Dissolve 5 g EDTA in 800 mL water and adjust the pH to 7.5 with NaOH. Stock solutions for all other components were prepared, filter sterilized and added in the amounts described below. Add water to 1 L and filter sterilize.

| Component | Stock concentration | Quantity for 1 L |
|---|---|---|
| $FeCl_3$ + 6H20 | | 830 mg |
| $ZnCl_2$ | | 84 mg |
| $CuCl_2 + 2H_2O$ | 1.7 g / 100 mL | 765 µL |
| $CoCl_2 + 2H_2O$ | 4.76 g / 100 mL | 210 µL |
| $H_3BO_3$ | 0.62 g / 100 mL | 1.6 mL |
| $MnCl_2 + 4H_2O$ | 19.8 g/ 100 mL | 8.1 µL |

**Table S6: Level 1 and level 2 plasmids assembled in this study.** Assembly was done within the framework of the Marburg Collection (Stukenberg *et al.*, 2021). Plasmid maps are provided in Supplementary Data 1.

| Plasmid name | Description |
|---|---|
| pST_236 | Level 1 plasmid with dCas9 transcription unit for construction of pST_300 and pST_301, Promoter: $P_{tet}$, RBS: B0029, CDS: dCas9, degradation tag: M0050, terminator: B0015 |
| pST_237 | Level 1 plasmid, mScarlet-I-HiBiT transcription unit, used as PCR template for preparation of tDNA |
| pST_240 | Empty plasmid as control |
| pST_300 | CRISPRi plasmid for single gRNAs |
| pST_301 | CRISPRi plasmid for multiple gRNAs |
| pST_302 | Level 1 plasmid, Fluc-HiBiT transcription unit, used as PCR template for preparation of tDNA |
| pST_303 | Level 1 plasmid, Rluc-HiBiT transcription unit, used as PCR template for preparation of tDNA |
| pST_310 | Level 1 plasmid, Nluc-HiBiT transcription unit, used as PCR template for preparation of tDNA |

**Table S7: *V. natriegens* strains with chromosomal modification**

| Strain | Genotype | Parental strain |
|---|---|---|
| DST016 | Δdns | V. natriegens ATCC14048 |
| DST018 | Δdns, int9::mScarlet-I-HiBiT | DST016 |
| DST023 | Δdns, argA-HiBiT | DST016 |
| DST024 | Δdns, metA-HiBiT | DST016 |
| DST025 | Δdns, eno-Hibit | DST016 |
| DST046 | Δdns, int12::Fluc-HiBiT | DST016 |
| DST047 | Δdns, int14::Rluc-HiBiT | DST016 |
| DST048 | Δdns, int9::mScarlet-HiBit, int12::Fluc-HiBit | DST018 |
| DST049 | Δdns, int9::mScarlet-HiBit, int12::Fluc-HiBiT, int14::Rluc-HiBiT | DST048 |
| DST050 | Δdns, int9::mScarlet-HiBit, int12::Fluc-HiBit, int14::Rluc-HiBit, int18::Nluc-HiBit | DST049 |
| DST051 | Δdns, int18::Nluc-HiBiT | DST016 |
| DST052 | Δdns, ΔargA | DST016 |
| DST053 | Δdns, ΔmetE | DST016 |

**Table S8: Recipe for SOC medium**

| Component | Concentration |
|---|---|
| Tryptone | 20 g/L |
| Yeast extract | 5 g/L |
| NaCl | 0.58 g/L |
| $H_20$ | Ad ~ 900 mL |
| Autoclave | |
| $MgSO_4$ (1 M) | 10 mL |
| $MgCl_2$ (1 M) | 10 mL |
| Glucose (2 M) | 10 mL |
| $H_20$ | Ad 1 L |

**Table S9: gRNA position vectors for assembly of multi gRNA CRISPRi plasmids.** Position of these gRNA cassettes according to the nomenclature of the Marburg Collection (Stukenberg *et al.*, 2021). Scheme for the assembly of multi gRNA CRISPRi plasmids is shown in Figure S15. Plasmid maps are provided in Supplementary Data 1.

| Plasmid name | Description |
|---|---|
| pMC0*_gRNA_Pos2_$P_{3b5b}$_pcaU (sacB-sfGFP) | Position 2, carries expression cassette for repressor *pcaU*, dropout *sacB-sfGFP* which is replaced by gRNA spacer |
| pMC0*_gRNA_Pos3_$P_{3b5b}$ (sacB-sfGFP) | Position 3, dropout *sacB-sfGFP* which is replaced by gRNA spacer |
| pMC0*_gRNA_Pos4_$P_{3b5b}$ (sacB-sfGFP) | Position 4, dropout *sacB-sfGFP* which is replaced by gRNA -spacer |
| pMC0*_gRNA_Pos5_$P_{3b5b}$ (sacB-sfGFP) | Position 5, dropout *sacB-sfGFP* which is replaced by gRNA spacer |
| pMC0*_gRNA_Pos3-5_$P_{3b5b}$ (sacB-sfGFP) | Position 3-5, dropout *sacB-sfGFP* which is replaced by gRNA spacer |
| pMC0*_gRNA_Pos4-5_$P_{3b5b}$ (sacB-sfGFP) | Position 4-5, dropout *sacB-sfGFP* which is replaced by gRNA spacer |

**Table S10: Protocol for microplate reader measurements without fluorescence measurement**

| | |
|---|---|
| Temperature | 37°C |
| Kinetic duration | 5 min |
| Shaking (orbital) duration | 120 s |
| Shaking (orbital) amplitude | 2 mm |
| Shaking (linear) duration | 120 s |
| Shaking (linear) amplitude | 2 mm |
| Absorbance measurement | |
| Measurement wavelength | 595 nm |
| Bandwidth | 10 nm |
| Number of flashes | 5 |
| Settle time | 10 ms |

**Table S11: Protocol for microplate reader measurements with fluorescence measurement**

| | |
|---|---|
| Temperature | 37°C |
| Kinetic duration | 5 min |
| Shaking (orbital) duration | 100 s |
| Shaking (orbital) amplitude | 2 mm |
| Shaking (linear) duration | 100 s |
| Shaking (linear) amplitude | 2 mm |
| Absorbance measurement | |
| Measurement wavelength | 595 nm |
| Bandwidth | 10 nm |
| Number of flashes | 5 |
| Settle time | 10 ms |
| Fluorescence measurement | |
| Excitation wavelength | 560 nm |
| Emission wavelength | 610 nm |
| Excitation bandwidth | 20 nm |
| Emission bandwidth | 20 nm |
| Gain | 50 |
| Number of flashes | 5 |
| Integration time | 20 µs |
| Lag time | 0 µs |
| Settle time | 10 ms |
| Mirror (automatic) | 50% mirror |
